# Key moments in naturalistic events synchronize neural activity patterns and dominate memory reinstatement

**DOI:** 10.1101/2025.08.30.673233

**Authors:** Aditya Upadhyayula, John M. Henderson, Jeffrey M. Zacks, Zachariah M. Reagh

**Affiliations:** Department of Psychological & Brain Sciences, Washington University at St. Louis.; Department of Psychology, University of California, Davis; Center for Mind and Brain, University of California, Davis

## Abstract

Continuous experiences are experienced and remembered in terms of events that unfold over time. There is strong evidence that event boundaries segment experience during comprehension and that event representations are compressed in memory; however, this compression is poorly understood. We developed a novel storyboard paradigm to test the hypothesis that representations of continuous experiences are defined by a subset of key moments that capture the underlying narrative. Participants agreed on when key moments occurred; some key moments corresponded with event boundaries, but many did not. fMRI during encoding revealed that neural activity patterns throughout the default network synchronized across individuals at key moments. Further, comparing fMRI during encoding to fMRI during retrieval revealed that key moments are overrepresented in neural patterns that are reinstated during event recall in the posterior-medial cortex. These results suggest that continuous events are punctuated by a small subset of meaningful moments, which dominate neural representations during perception and memory.

## Introduction

Not all moments in life are equal. As individuals move through continuous streams of experience, some moments are perceived as being especially meaningful or memorable. For instance, when reflecting on a movie, one might recall just a few pivotal scenes — a shocking twist, an emotional climax, or the final resolution — while the rest fades into the background. These moments tend to anchor one’s understanding of the story—a potentially efficient form of compression in memory. Despite the importance of such *key moments* for comprehension and memory, their cognitive and neural mechanisms are poorly understood.

Furthermore, behavioral^1–3^ and neural evidence^4–6^ indicates that continuous experiences are processed and remembered as discrete events. Regions of the brain’s Default Mode Network (DMN) are increasingly thought to be critical for this process. For example, when people follow the same story, patterns of activity in the DMN regions synchronize across participants^7–10^. These regions don’t just keep track of the story as it unfolds – they represent individual events^11^ and their structure^5^, and carry knowledge about event schemas^12^, and the individual components that make up the events^13^. *Event boundaries* – moments when one event ends and another begins – serve to scaffold the external structure of events by delineating ongoing experience into two units. This process triggers spikes of activity in the hippocampus^6^ and elicits pattern shifts in the brain networks (especially DMN) that help reorganize how ongoing situations are represented^4–6^. In contrast to the decades of research underscoring the role of event boundaries in shaping event understanding and memory^14–16^, little is known about what governs the structure of key moments within events^17,18^. How does the brain encode and retrieve information about particularly important moments that may or may not be boundaries?

Understanding the structure of events is crucial to better understand memory. Memory is inherently lossy and favors abstraction and compression over time, preserving high-level event structure and meaning^11,12,19^. This transformation is also evident from the literature on aging, where information about high-level event schemas is preserved compared to the low-level perceptual features^20,21^, and also in memory replay, where the temporal structure of past experiences is compressed^22,23^. Such compression implies that the brain does not reactivate every moment but may instead rely on a subset of informative or representative moments to scaffold recall. However, the format and content of this compression—and the mechanism by which the brain selects specific moments within an event for reinstatement—remain poorly understood. In this study, we explore key moments to investigate the structure of events and their role in event recall.

The concept of key moments is deeply familiar in everyday life. When recounting the vacation experience to a friend, people don’t narrate every step. Instead, a handful of pivotal moments are often highlighted: the breathtaking view, the missed train, the big family dinner. Similarly, educational tools like CliffsNotes and flashcards condense broader narratives or topics into the most essential elements, emphasizing representative ideas or key points to support efficient learning and memory. Comics are a predominant media form that uses key moments in storytelling^26,27^. Film makers formalize this process in commercial Hollywood films^24,25^ with storyboards, which outline the story through a sequence of carefully chosen set of still frames. Across these domains, the emphasis is on a representative set of moments that carry meaning, organize the experience, and make it memorable. Yet despite how central this principle is in daily life, the scientific study of which moments emerge as “key” and why remains limited.

What constitutes a key moment? One possibility is that event boundaries themselves are key moments, since they tend to occur at moments of substantial change – such as a shift in location, time, or action – that signal the transition from one event to another. Indeed, early work by Newtson and colleagues^3,28^ found that deleting information from event boundaries impaired comprehension and memory more than deleting equivalent amounts of information from event middles. Subsequent investigations in event cognition have generally found that event middles are less well remembered than moments around event boundaries^29^. However, not all moments within events are equally meaningful; some carry particular importance due to their emotional weight, narrative significance, or information richness^17,30^. Thus, key moments may not be limited to transitions between events, but could also include especially salient moments within them.

Here, we tested the hypotheses that continuous experiences are represented in part as a series of key moments, which are somewhat distinct from event boundaries, and that these moments are essential components of memory representations. We developed a novel storyboard paradigm to identify these key moments normatively (Figure 1). Participants watched movie clips and were subsequently instructed to communicate the underlying story by selecting the individual image frames that made up the story in each clip. People showed strong agreement in which frames were selected, and importantly, these key moments were distinct from event boundaries. Analysis of an open fMRI dataset where participants viewed the same movie revealed that neural activity patterns synchronized strongly across individuals at key moments throughout the Default Mode Network (DMN). Further, as participants recalled the movie, neural patterns associated with key moments dominated the patterns reinstated in the posterior-medial cortex. These findings demonstrate that event representations are characterized by a relatively small subset of meaningful moments.

**Figure 1:**
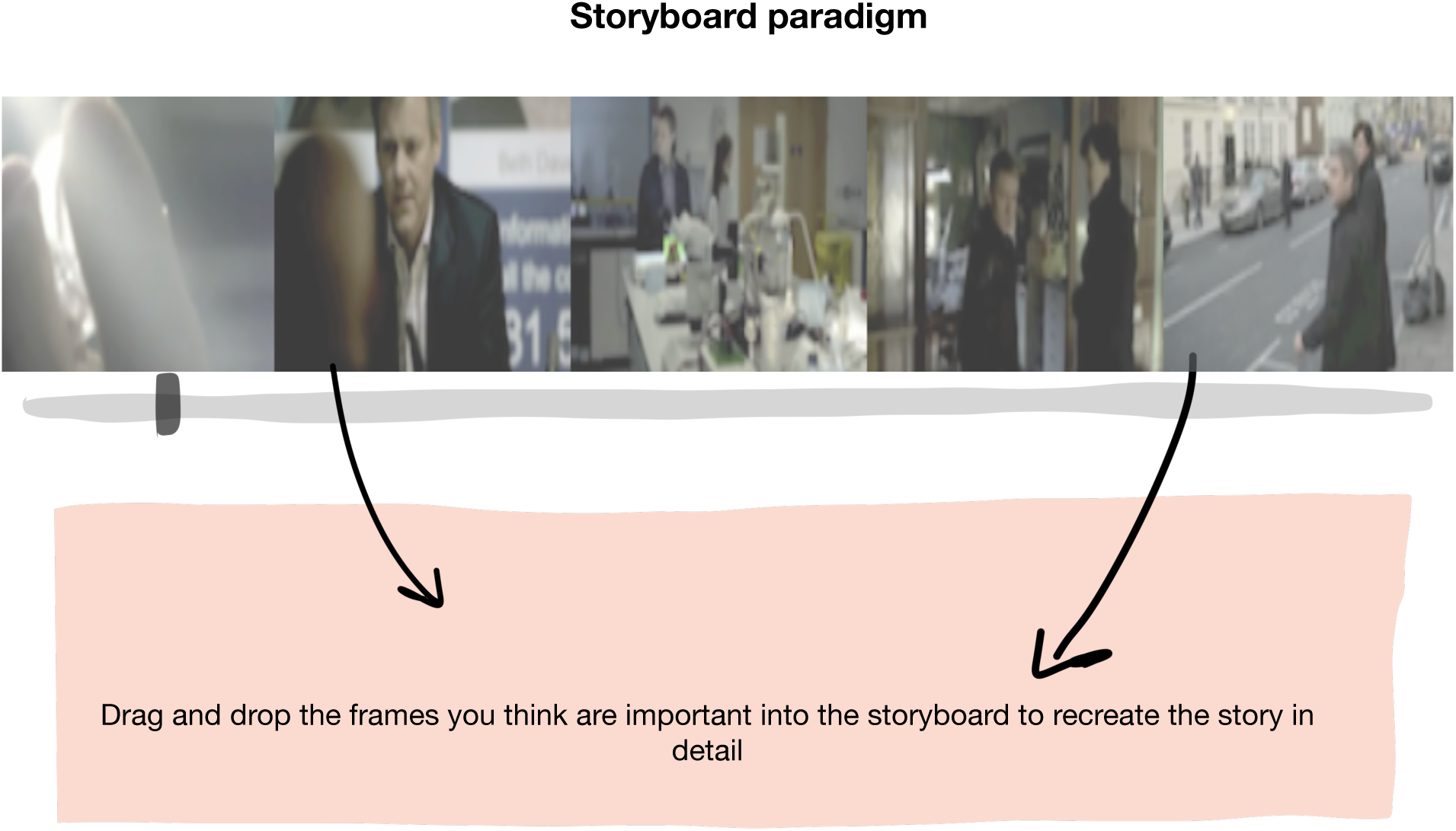
Schematic illustrating the storyboard paradigm to identify key moments in a movie. Participants were first shown a movie without any interruption and were then presented with the individual frames that make up the entire movie. They were instructed to pick out the frames they thought were important to recreate the story by dragging and dropping such frames into the storyboard.

## Results

### Participants agree on key moments in a movie, which are partially distinct from event boundaries

One hundred ninety-eight participants from an online sample viewed movie clips, separated at experimenter-identified coarse-grained event boundaries (ranging from 11- 185 seconds long), from BBC’s Sherlock (S1E1: *A Study in Pink*). After viewing each clip, participants completed the storyboard task (Figure 1). Each segmented clip was split into individual frames, which the participants used to construct the underlying story. Participants were instructed to recreate the story in detail using the still images provided (i.e., selecting key moments), and were not given any minimum or maximum number of frames to select. During selection, participants had access to buttons to Undo, Redo, Reorder, and Clear their storyboards. After exclusions for internet connectivity issues or lack of task understanding, 189 participants were retained, with at least 45 participants providing responses for a given clip. We aimed to test whether key moments captured aspects of continuous experiences that were distinct from event boundaries. For comparisons with event boundaries, we used segmentation data gathered by Chen & Swallow (2024)^31^. Briefly, for segmentation, 39 participants provided measures of both coarse and fine-grained event boundaries for all the clips within the same Sherlock episode.

On average, participants identified 9.6 key moments (SD = 3.9) and 8.9 fine-grained event boundaries (SD = 5.9) per clip. Thus, people showed a similar timescale of selecting key moments and segmenting events. For both key moments and event boundaries, we convolved the time-series of key moments with a Gaussian distribution (FWHM ≈ 1.5 sec) to account for noise in the individual responses. We next constructed leave-one-out population distributions for both key moments and event boundaries by averaging the Gaussian convolved participant responses. To assess how well each individual aligned with the group, the correlation between individual distribution and the corresponding leave-one-out average was computed. Furthermore, to test against chance, each individual distribution was then compared against a permuted null distribution obtained by randomly selecting the key moments or event boundaries while preserving the interval durations (Figure 2A; see Methods).

**Figure 2:**
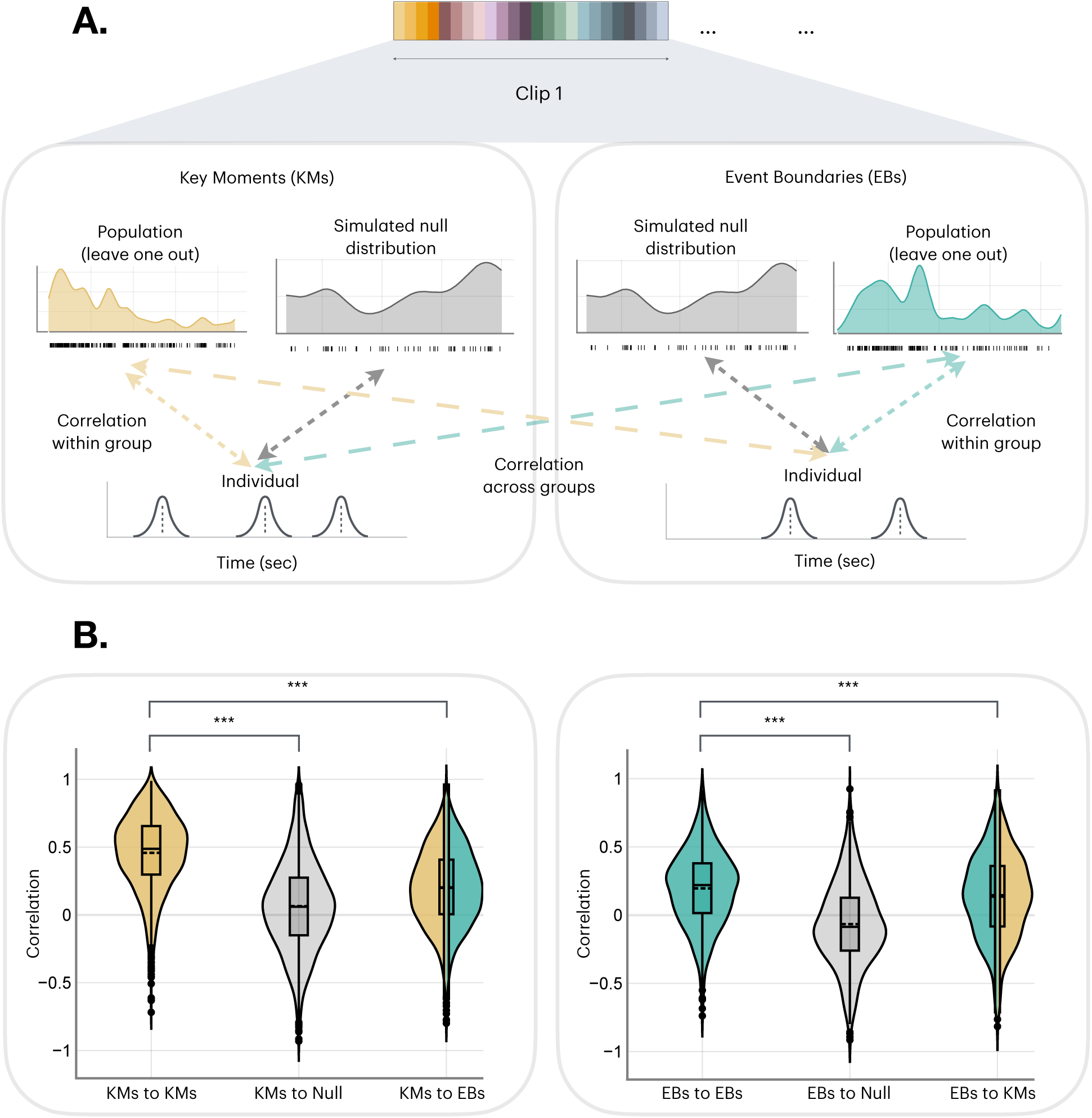
A. Schematic illustrating the correlation analyses comparing Key Moments (KMs) to Event Boundaries (EBs) and simulated null distributions. Individual distributions from each clip (as depicted by the rainbow bar above) are correlated with the leave-one-out average population distribution (yellow and green color arrows within the two respective boxes in the top panel), a randomly sampled null distribution (the gray arrows in both boxes of the top panel), and the cross-population average distribution (the yellow and green colored arrows between the two boxes in the top panel). The obtained correlations are then compared using linear mixed-effects models to determine if there are systematic differences between the KM and EB distributions. B. Left panel: The yellow violin plot shows the distribution of correlations between individual KM and the KM population (leave-one-out) average distributions. The gray violin plot shows the distribution of correlations between individual KM distributions and simulated null distributions. The mixed color violin plot shows the distribution of correlations between individual KM distributions and the population average of the EB distributions. The boxes in the violin plots represent the 25th and 75th quartile ranges. Solid lines describe the median, and the dashed line represents the mean of the distribution. Right panel: Same analysis as the left panel, except EB distributions are compared to a randomly sampled null distribution and to the KM average distribution. Significance markers indicate p < 0.001

Results from linear mixed effects models indicated that with minimal instruction and no practice participants strongly agreed with each other on the key moments: The observed correlations far exceeded the mean of a null, shuffled key moment distribution (β = 0.40, SEM = 0.03, t(53.5) = 12.627, p < 0.001, Figure 2B). Thus, participants strongly agreed with each other on which frames were indicated to be key moments in the storyboard task. Participants also strongly agreed on event boundaries, exceeding a null distribution (β = 0.25, SE = 0.03, *t*(26.72) = 7.809, *p* < 0.001) – replicating prior work^2,32^. However, when comparing each participant’s key moment or event boundary distributions to the average of the other distribution, we found significantly higher within than across-task agreement (key moments vs. event boundary distribution: β = 0.25, SEM = 0.01, *t*(45.2) = 13.361, *p* < 0.001) (event boundaries vs. key moment distribution: β = 0.05, SEM = 0.01, *t*(44.45) = 3.632, *p* = 0.0007, Figure 2B). Finally, the cross-task agreements were also significantly different from zero (KM to EBs vs 0: mean = 0.209, SE = 0.02, t(51.98) = 9.095, p < 0.001 ), and (EBs to KMs vs. 0: mean = 0.13, SE = 0.02, t(44.9) = 5.881, p < 0.001), suggesting that although key moments and event boundaries differ from one another, they are correlated. Overall, these results suggest that the distribution of key moments does not fully overlap with event boundaries.

### Neural activity patterns synchronize across people at key moments

Brain activity synchronizes across people as they encode the same movie or audio narrative^7,9,11^, which is thought to reflect the shared understanding of the stimulus^7,11^. Here, we applied data from our storyboard paradigm to an fMRI dataset during viewing and recall of the same Sherlock episode (n=17), testing the hypothesis that key moments effectively summarize and anchor people’s shared representations of continuous events. We computed Inter-Subject neural Pattern Similarity (ISPS) to measure the multi-voxel pattern correlation of BOLD activity across participants as the movie unfolded. This was done by correlating the activity pattern for a given subject and TR within a clip with the leave-one-out average activity pattern from the other 16 subjects for the same TR. To test the influence of key moments in driving pattern similarity across people, we ran a linear mixed effects model predicting ISPS, with key moment and event boundary distribution probabilities as predictors (see Figure 3 for a schematic of the analysis; also see fMRI methods for more information aligning the granularity of both key moment and event boundary distributions to the neural data). This was repeated for all 46 clips for which the storyboard and fine-grained segmentation probability density distributions were obtained.

**Figure 3:**
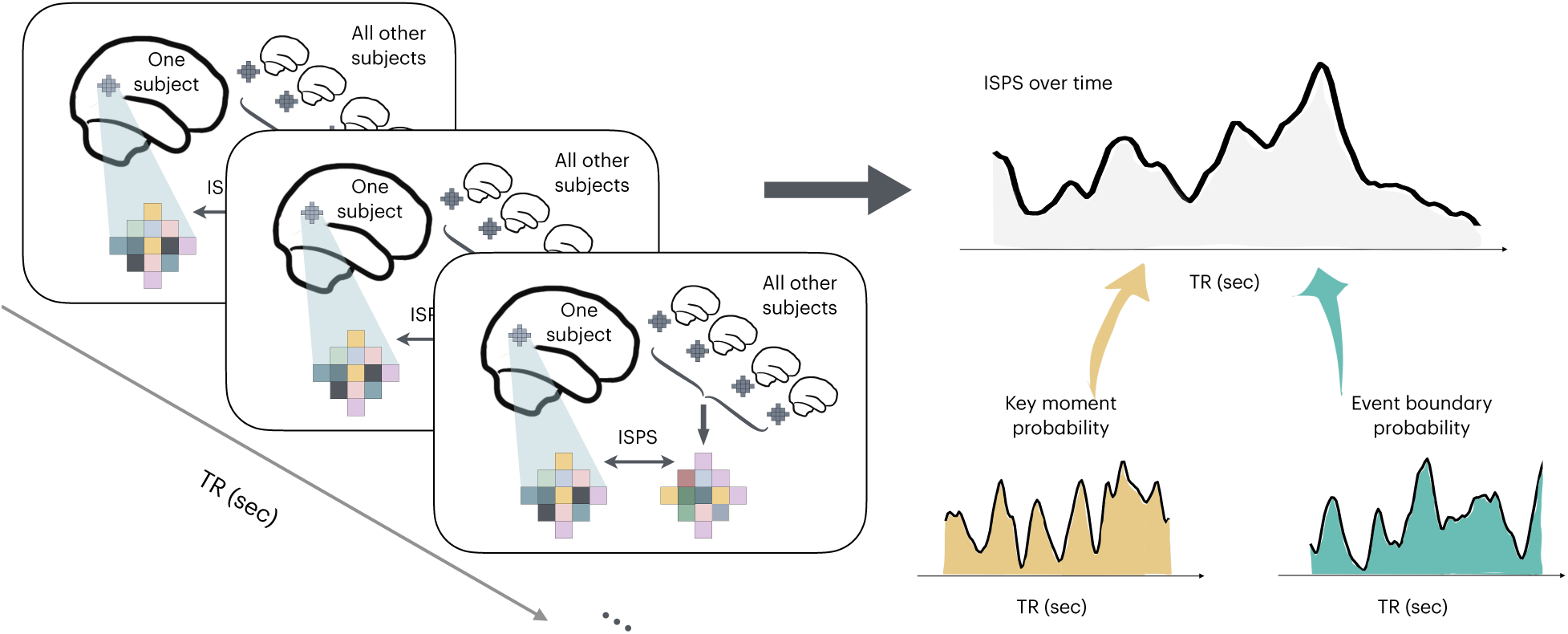
Schematic of the Intersubject Neural Pattern Synchrony (ISPS) computation. For each clip, a subject’s pattern from an ROI at a time point is compared to the rest of the subjects’ average pattern for that time point. The resulting ISPS signal is derived for all possible time points as measured in TRs, then compared against the key moment and the fine-grained event boundary probability distributions over TRs to see which of the two distributions captured the ISPS signal better.

Regions of Interest (ROIs) used in this analysis largely focused on the regions within the DMN highlighted in the prior studies of event representation and memory^11,13,21,33–35^. These included the posterior medial cortex (PMC), hippocampus (Hipp), parahippocampal cortex (PHC), medial prefrontal cortex (mPFC), and the perirhinal cortex (PRC). In addition, the early visual cortex (EVC) and the early auditory cortex (EAC) were also used as control regions in the analyses.

Before examining the influence of key moments and event boundaries on ISPS, we tested the reliability of the ISPS signal within our ROIs. We did this by testing whether the true ISPS exceeded each participant’s correlation with a null distribution ISPS obtained from the flipped average time series across other participants for the same clip. These true vs. null comparisons were significantly different across all ROIs (Table 1; Figure 4), indicating reliable ISPS effects across all regions examined.

**Figure 4:**
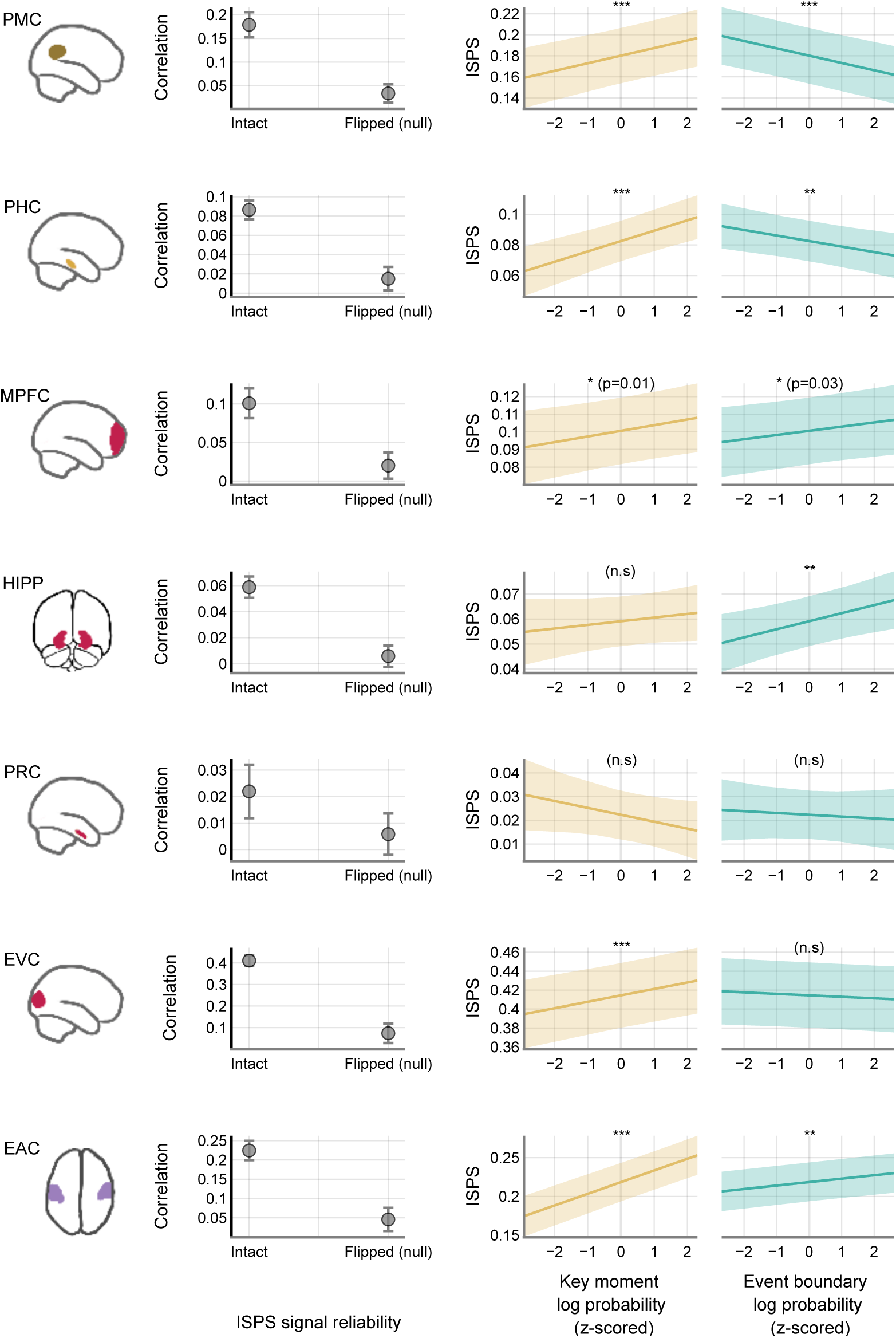
Comparing Inter-Subject neural Pattern Similarity (ISPS) against the key moment and event boundary probabilities. The first column indicates the ROI that was examined. The second column shows the reliability of the ISPS signal by comparing the ISPS signals against their flipped null distribution counterparts. The third column plots the relationship between the ISPS and the key moment probability distribution after log-transformed and z-scored. The fourth column shows the relationship between ISPS and the event boundary probability. Error bars and the shaded regions depict the 95% CI. The asterisks *** indicate a significance value of < 0.0001, ** indicate a significance value of < 0.01, and (n.s) indicates no significance.

**Table 1:**
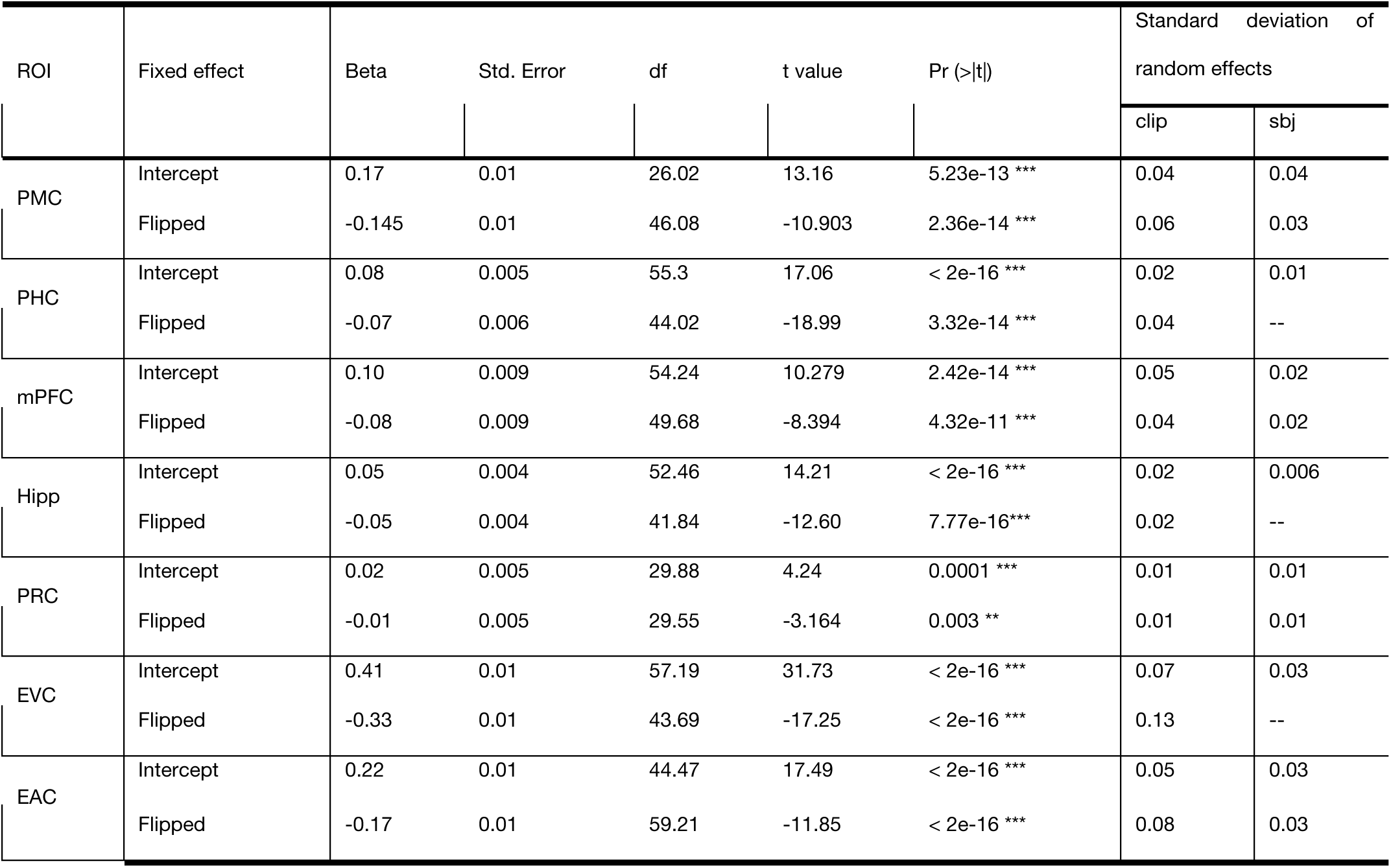
Linear Mixed Effects models testing the reliability of the measured ISPS signals across various ROIs. S.D is the standard deviation of the random effects.

The results from a linear mixed effects model showed a significant positive correlation between the key moment probabilities and the ISPS scores in the PMC (β = 0.006, SEM = 0.002, df = 2.351e4, t = 3.325, p = 0.0008), a region previously found to carry event-specific neural patterns^5,11,33,34^ (Figure 4). We also found positive relationships between key moments and ISPS in the PHC (β = 0.007, SEM = 0.002, df = 1.586e4, t = 4.665, p < 0.0001), mPFC (β = 0.003, SEM = 0.001, df = 2.943e4, t = 2.439, p = 0.0188), as well as EVC (β = 0.006, SEM = 0.002, df = 2.848e4, t = 3.688, p = 0.0002) and EAC (β = 0.015, SEM = 0.001, df = 2.975e4, t = 10.608, p < 0.0001). Thus, across many of our *a priori* ROIs, the probability that a given moment was selected as a key moment is associated with stronger ISPS. On the other hand, event boundaries had a significant negative effect on the ISPS in PMC (β = -0.006, SEM = 0.001, df = 3.126e4, t =-4.807, p < 0.0001) and PHC (β = -0.003, SEM = 0.001, df = 3.089e4, t = -2.927, p = 0.003), and were positively associated with ISPS in the mPFC (β = 0.002, SEM = 0.001, df = 3.136e4, t =2.119, p = 0.0341) and the hippocampus (HPC) (β = 0.003, SEM = 0.001, df = 3.081e4, t =2.832, p = 0.004). There was no effect of key moments and event boundaries on the ISPS in perirhinal cortex (PRC) (KM: β = -0.002, SEM = 0.001, df = 2.996e3, t =-1.647, p = 0.099; EB: β = -0.0007, SEM = 0.001, df = 2.746e4, t =-0.5, p = 0.61) (see Figure 4; Table 2). Moreover, the interaction between event boundaries and key moments was also a significant predictor of the ISPS in the PMC, PHC, EVC, and EAC regions.

**Table 2:** Linear mixed-effects models comparing the ISPS activity over time with key moments and event boundary probability distributions.

| ROI | Fixed effect | Beta | Std. Error | df | t value | Pr (>\|t\|) | Standard deviation of random effects |  |
| --- | --- | --- | --- | --- | --- | --- | --- | --- |
|  |  |  |  |  |  |  | clip | sbj |
| PMC | Intercept | 0.18 | 0.013 | 25.83 | 13.462 | < 0.0001 *** | 0.04 | 0.04 |
|  | Key Moment | 0.007 | 0.002 | 2.106e4 | 4.129 | < 0.0001 *** | -- | -- |
|  | Event boundary | -0.006 | 0.001 | 3.126e4 | -4.807 | < 0.0001 *** | -- | -- |
|  | Interaction | -0.003 | 0.001 | 3.104e4 | -2.432 | 0.015 * | -- | -- |
| PHC | Intercept | 0.08 | 0.006 | 35.69 | 12.323 | < 0.0001 *** | 0.02 | 0.02 |
|  | Key Moment | 0.006 | 0.001 | 1.296e4 | 4.582 | 4.66e-06 *** | -- | -- |
|  | Event boundary | -0.003 | 0.001 | 3.089e4 | -2.927 | 0.00342 ** | -- | -- |
|  | Interaction | 0.006 | 0.001 | 3.028e4 | 5.611 | 2.03e-08 *** | -- | -- |
| mPFC | Intercept | 0.1 | 0.009 | 53.97 | 10.486 | < 0.0001 *** | 0.04 | 0.02 |
|  | Key Moment | 0.003 | 0.001 | 2.870e4 | 2.349 | 0.0188 * | -- | -- |
|  | Event boundary | 0.002 | 0.001 | 3.136e4 | 2.119 | 0.0341 * | -- | -- |
|  | Interaction | -0.004 | 0.001 | 3.134e4 | -0.405 | 0.6855 | -- | -- |
| Hipp | Intercept | 0.05 | 0.005 | 48.15 | 11.718 | 1.05e-15 *** | 0.02 | 0.01 |
|  | Key Moment | 0.001 | 0.001 | 1.231e4 | 1.070 | 0.28468 | -- | -- |
|  | Event boundary | 0.003 | 0.001 | 3.081e4 | 2.832 | 0.00463 ** | -- | -- |
|  | Interaction | -0.0009 | 0.001 | 3.014e4 | -0.838 | 0.40206 | -- | -- |
| PRC | Intercept | 0.02 | 0.005 | 29.84 | 4.316 | 0.0001 *** | 0.01 | 0.01 |
|  | Key Moment | -0.002 | 0.0017 | 2.996e3 | -1.647 | 0.0996 | -- | -- |
|  | Event boundary | -7.6e-4 | 0.001 | 2.746e4 | -0.50 | 0.616 | -- | -- |
|  | Interaction | 0.0003 | 0.001 | 2.513e4 | 0.235 | 0.8138 | -- | -- |
| EVC | Intercept | 0.41 | 0.01 | 32.33 | 23.897 | < 2e-16 *** | 0.06 | 0.05 |
|  | Key Moment | 0.006 | 0.002 | 2.848e4 | 3.688 | 0.0002 *** | -- | -- |
|  | Event boundary | -0.001 | 0.001 | 3.117e4 | -1.043 | 0.296 | -- | -- |
|  | Interaction | 0.003 | 0.001 | 3.114e4 | 2.089 | 0.03 * | -- | -- |
| EAC | Intercept | 0.21 | 0.01 | 44.29 | 17.421 | < 2e-16 *** | 0.05 | 0.03 |
|  | Key Moment | 0.015 | 0.001 | 2.975e4 | 10.608 | < 2e-16 *** | -- | -- |
|  | Event boundary | 0.004 | 0.001 | 3.138e4 | 3.825 | 0.000131 *** | -- | -- |
|  | Interaction | 0.005 | 0.001 | 3.137e4 | 4.673 | 2.98e-6 *** | -- | -- |

As an alternative, we looked at a permutation analysis comparing shuffled key moment and event boundary distributions against the intact distributions to predict the ISPS signal. We did this analysis because the previous analysis could be susceptible to inflation of effects due to temporal autocorrelation in the large number of time points. This analysis revealed that the effect of intact key moment and event boundary distributions was significantly different from their shuffled counterparts in predicting the ISPS signal, and the effects were in line with what we observed above (see Supplementary materials S1 - Figure S 1 for more information). Finally, we also ran an additional analysis where we separated the key moment and event boundary probability distributions into shared and unique components of each. The unique KM and EB probability distributions were then used to predict the ISPS (see Supplementary materials S4 - Figure S 5 for more information). This analysis also replicated the findings in the above analyses. Overall, these results suggest that intersubject neural synchrony is particularly strong at key moments in PMC and other regions. Furthermore, this effect is distinct from the effect of event boundaries on ISPS.

### Key moments dominate neural pattern reinstatement during event recall

Reliable spatial reactivation of encoding patterns during recall has been reported across the DMN^11,19,36,37^ for naturalistic stimuli. These studies have typically correlated brain activation patterns during movie viewing with those of spoken recall, either at the whole-event level^11,36^ or at the subject level^19^. These encoding-retrieval pattern similarity analyses are often constrained for two reasons: First, not every stimulus will be recalled by the participants. Second, recall durations vary across people and are typically shorter than the actual duration of the stimulus, complicating comparisons with encoding with specificity beyond whole events. Existing analyses^11,13^ have largely compared neural activity averaged across all recall timepoints against activity averaged across all encoding timepoints to overcome this challenge. However, key moments pertain to individual events during encoding. Therefore, understanding their effects on recall at a finer grain would be useful to better understand neural representation of events. Before focusing on the effects of key moments on reinstatement patterns during recall, which depends on moment-to-moment movie viewing and recall comparisons of the neural data, we first wanted to establish whether individual moments during movie-viewing reliably correlated with the average neural pattern present during event recall. We validated this approach by correlating each movie-viewing TR’s neural pattern with the average recall pattern of the same clip and participant (referred to as within-clip condition). As a null comparison, each movie-viewing TR’s neural pattern was compared to the average recall pattern from the rest of the clips and the same subject (across-clips condition). This analysis revealed a significant difference between the correlation distributions for within-clip and across-clips conditions in the PMC (β = -0.145, SEM = 0.01, df = 46.08, t =-10.903, p < 0.0001), mPFC (β = 0.02, SEM = 0.01, df = 46.08, t =-10.903, p < 0.0001), and the PHC (β = 0.01, SEM = 0.006, df = 29.38, t =-2.323, p =0.02) (Figure 5, Table 3). That is, recall patterns PMC, PHC, and MPFC reliably carried information about individual moments present during encoding.

**Figure 5:**
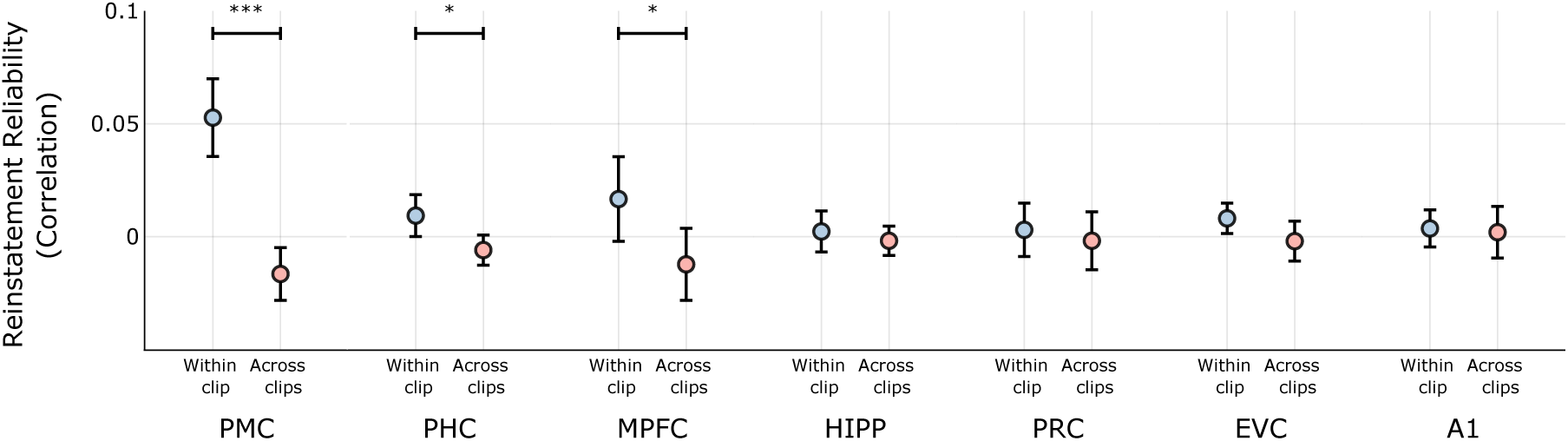
Moment-to-moment reinstatement reliability across various ROIs. Each TR’s activity pattern was compared with the average recall pattern for that clip (within-clip), and also with the average recall activity patterns of the remainder of the 45 clips (across-clips). The correlations were then compared using linear mixed-effects models suggesting a significant effect between the conditions across the Posterior Medial Cortex (PMC), Parahippocampal Cortex (PHC), and the medial Prefrontal Cortex (mPFC). Individual dots represent the model estimates, and the error bars reflect the 95% CI of the model estimates.

**Table 3:**
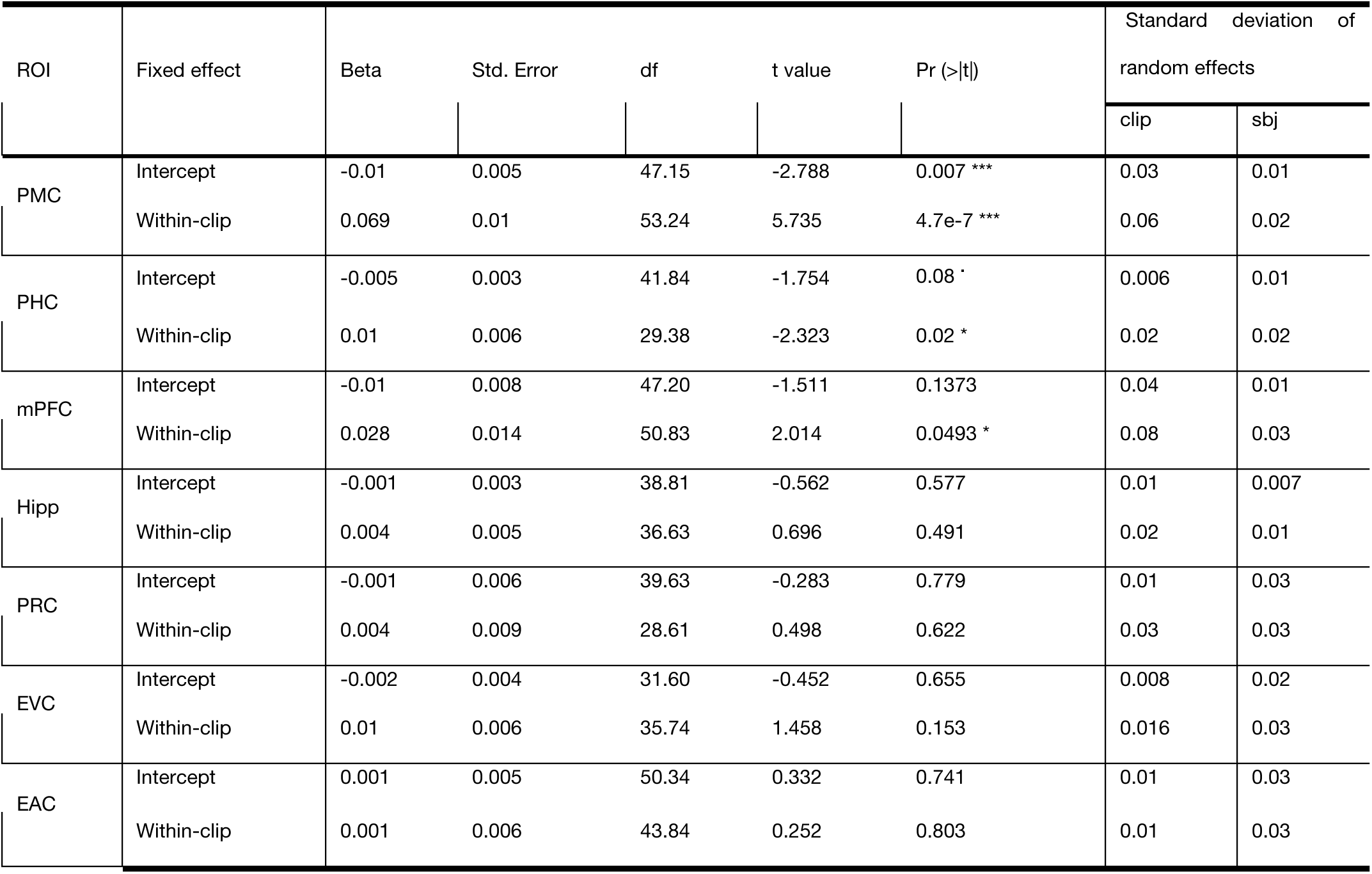
Linear mixed effects models evaluating Moment-to-moment reinstatement reliability (Neural Pattern Reinstatement) between encoding and recall across various Regions of Interest.

We next analyzed whether the patterns at key moments and at event boundaries uniquely predicted recall activation patterns within these ROIs. If key moments are important for how the experience is remembered, then the neural patterns corresponding to these specific moments during encoding should be reflected in the neural activity present during recall. To test this hypothesis, we compared the average recall pattern directly to the movie-viewing pattern at each TR, creating a measure of neural pattern reinstatement (NPR) that is time-resolved across encoding patterns. Our critical question whether key moments, event boundaries, or both showed uniquely strong NPR. See Figure 6A for a schematic outlining the analysis. A linear mixed effects model analysis was implemented with the NPR as the dependent variable, and storyboard and event boundary probabilities as the predictor variables. Subjects and clips were used as random effects with clips nested within subjects. This analysis revealed a significant main effect of the key moment probabilities (beta = 0.005, SE = 0.002, t(2.08e4) = 2.554, p = 0.01) in predicting pattern reinstatement within the PMC (Figure 6B). However, event boundary probabilities were not significant predictors of pattern reinstatement in this region (β = -0.001, SE = 0.0017, t(246) = -1.008, p = 0.313). These effects were not observed in the mPFC – (key moments: β = 0.002, SE = 0.001, t(2.48e4) = -0.183, p = 0.237; event boundaries: β = -0.001, SE = 0.001, t(2.45e4) = -0.81, p = 0.418) or PHC - key moments: β = 0.0015, SE = 0.0016, t(1.78e4) = 0.951, p = 0.342; event boundaries: β = -0.0012, SE = 0.0014, t(2.49e4) = -0.858, p = 0.391) (Figure 6B; Table 4).

**Figure 6:**
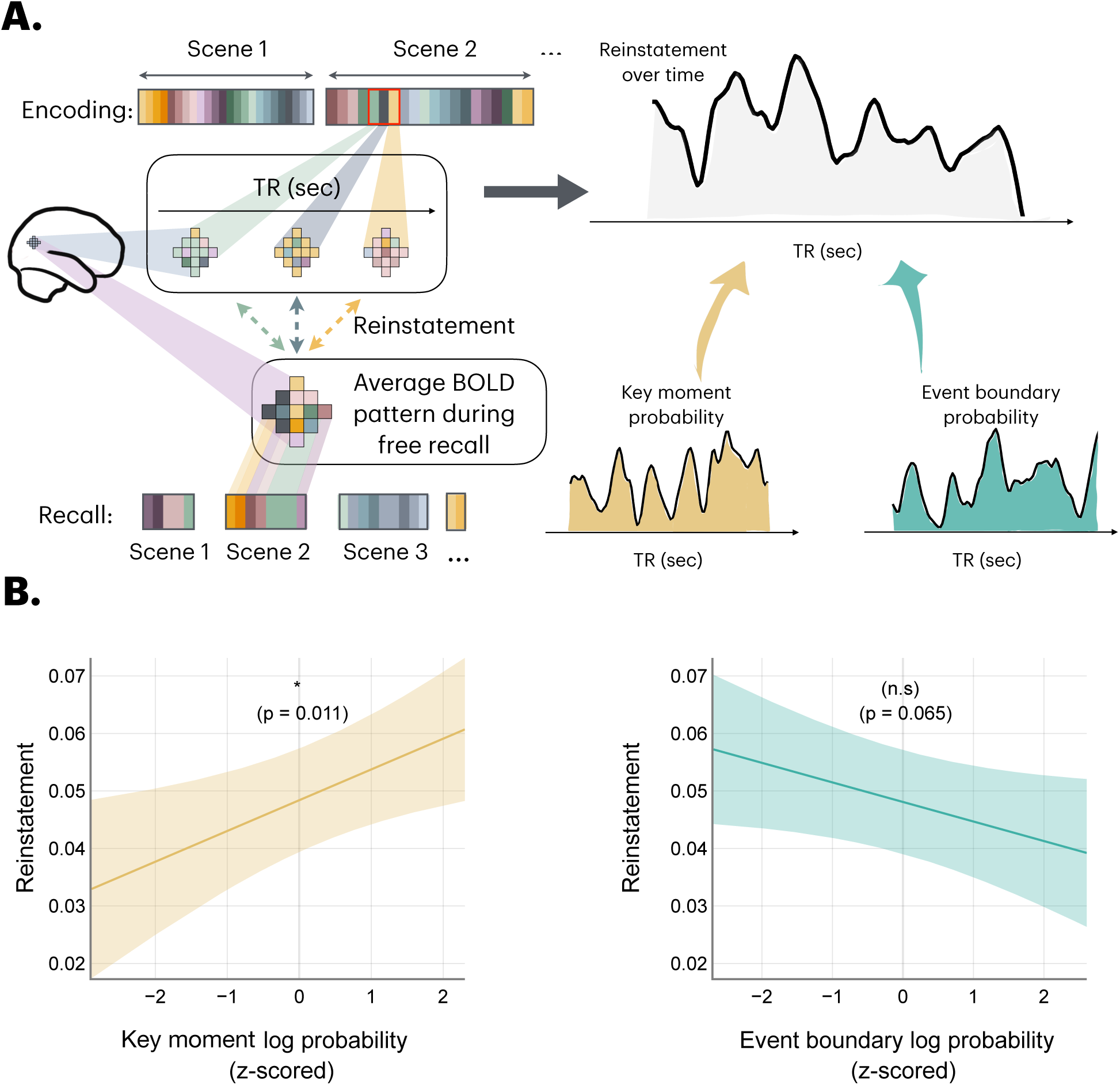
A). Schematic comparing the Neural Pattern Reinstatement over time against the storyboard and event boundary probabilities. For each successfully recalled clip, the average neural data of the recall was correlated with the moment-to-moment neural pattern present during movie viewing to create a Neural Pattern Reinstatement (NPR) over time. To see which of the moments during movie viewing were important for reinstatement, the NPR signal is then compared against the storyboard and event boundary probabilities for the given clip. B). Neural Reinstatement Pattern as a function of the Storyboard and Event boundary probability. Each probability distribution was log-transformed and z-scored to facilitate the linear mixed effects model comparison. TRs with a high probability of being key moments, as measured by the storyboard paradigm, were significant predictors of the NPR. However, event boundary probabilities were not significant predictors of the NPR signal. Lines indicate the beta estimate from the linear mixed effects models. Shaded regions indicate the 95% CI.

**Table 4:**
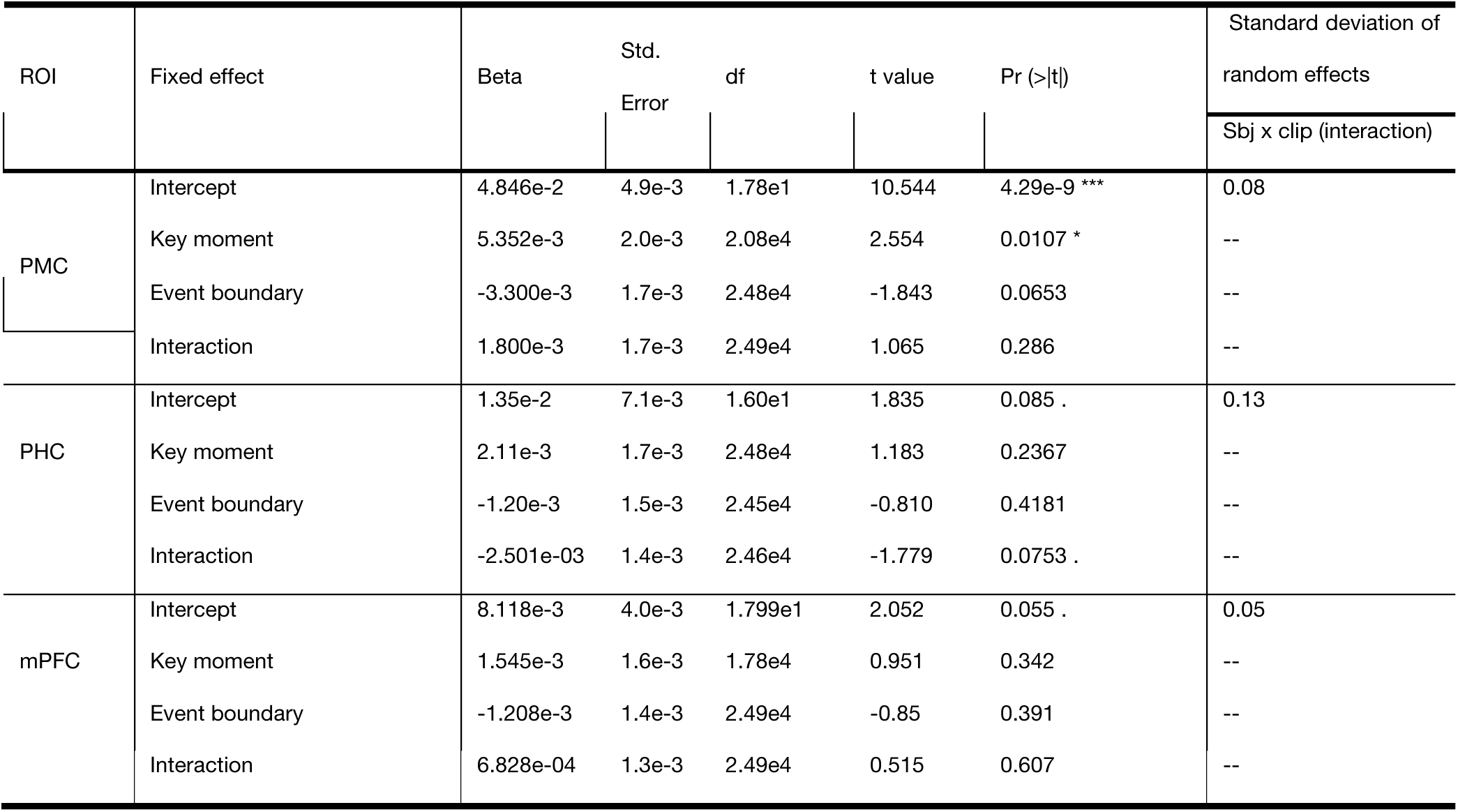
Linear Mixed Effects models evaluating the Neural Pattern Reinstatement (NPR) signal as a function of the log transformed and z-scored key moment and event boundary probabilities. Regions of Interest include Posterior Medial Cortex (PMC), the Posterior Hippocampal Cortex (PHC), and the medial Prefrontal Cortex (mPFC). These regions of interest were chosen following the NPR reliability analysis (see Table 3 for more information).

| ROI | Fixed effect | Beta | Std.<br>Error | df | t value | Pr (>\|t\|) | Standard deviation of<br>random effects |
| --- | --- | --- | --- | --- | --- | --- | --- |
|  |  |  |  |  |  |  | Sbj x clip (interaction) |
| PMC | Intercept | 4.846e-2 | 4.9e-3 | 1.78e1 | 10.544 | 4.29e-9 *** | 0.08 |
|  | Key moment | 5.352e-3 | 2.0e-3 | 2.08e4 | 2.554 | 0.0107 * | -- |
|  | Event boundary | -3.300e-3 | 1.7e-3 | 2.48e4 | -1.843 | 0.0653 | -- |
|  | Interaction | 1.800e-3 | 1.7e-3 | 2.49e4 | 1.065 | 0.286 | -- |
| PHC | Intercept | 1.35e-2 | 7.1e-3 | 1.60e1 | 1.835 | 0.085 . | 0.13 |
|  | Key moment | 2.11e-3 | 1.7e-3 | 2.48e4 | 1.183 | 0.2367 | -- |
|  | Event boundary | -1.20e-3 | 1.5e-3 | 2.45e4 | -0.810 | 0.4181 | -- |
|  | Interaction | -2.501e-03 | 1.4e-3 | 2.46e4 | -1.779 | 0.0753 . | -- |
| mPFC | Intercept | 8.118e-3 | 4.0e-3 | 1.799e1 | 2.052 | 0.055 . | 0.05 |
|  | Key moment | 1.545e-3 | 1.6e-3 | 1.78e4 | 0.951 | 0.342 | -- |
|  | Event boundary | -1.208e-3 | 1.4e-3 | 2.49e4 | -0.85 | 0.391 | -- |
|  | Interaction | 6.828e-04 | 1.3e-3 | 2.49e4 | 0.515 | 0.607 | -- |

In addition to the above analysis, we performed an analysis in which we computed weighted patterns during encoding, with weighting being determined by key moment or event boundary probability, and compared them to event-averaged recall patterns. This approach is similar to prior whole-event encoding-recall comparisons used in the literature ^11,13^ (see Supplemental analysis S2 for more information). Briefly, this analysis converged with the TR-by-TR analysis in that the key-moment weighted movie-viewing encoding activity patterns were over-represented in the recall neural activity patterns in PMC.

## Discussion

This study investigated the hypothesis that continuous experiences are represented, in part, as key moments that anchor ongoing event representations and are central to event memories. We found that people reliably picked certain moments as important for conveying the underlying story. Both behavioral and neural analyses revealed that these key moments capture distinct aspects of the experience that are partially dissociable from event boundaries. In a subset of posterior regions of the DMN (PMC and PHC), participants’ brain activity patterns during movie viewing were more similar to each other at these key moments, unlike event boundaries, where the brain activity patterns were desynchronized. Notably, an opposite profile was found for an anterior DMN region (mPFC) as well as HPC, where the brain activity patterns during movie viewing were more synchronized during event boundaries. The DMN has been implicated in tracking high-level information such as context and situation during comprehension of complex experiences^9–11,13,19,33,35,38^, and the sensitivity of the sub-regions within this network to these key moments suggests that they capture crucial elements of narrative comprehension. Furthermore, within the PMC and the mPFC, the neural patterns corresponding to these key moments were reinstated during the recall of the corresponding scenes. The fact that the PHC of the DMN were more sensitive to the key moments, whereas the mPFC and hippocampus were sensitive to the event boundaries, suggests that specialized mechanisms regulate the encoding of key moments into memory and the reconstruction of a scene from those moments during subsequent recall.

Previous work has focused on the segmentation of the stream of behavior into discrete units. Less is known about how those units are internally structured. Previous studies suggest that, overall, event middles are less well remembered than are boundaries^3,28,39,40^. Following proposals by Newtson^3,28,40^ Event Segmentation Theory (EST)^41^ proposed that event boundaries form the scaffold of subsequent memory. Event boundaries are generally moments of distinctiveness^14,41–44^, which is an adaptive attribute for a memory scaffold. However, event boundaries also are generally moments of high feature change as measured by neural pattern shifts^12^, which may not be well suited to representing stable configurations of features that are important for comprehension; thus, comprehending or recalling an event might therefore depend not only on that event boundaries, but also on specific key moments that serve to anchor or summarize the representation.

Further, it is also possible that the relationship between key moments and event boundaries may vary depending on the event type. Consider a coffee-making scenario involving pouring coffee into a mug. Intuitively, the most important and meaningful moments might seem to be at the event’s beginning and end – where the coffee mug is placed under the coffee machine, and where someone ultimately retrieves their coffee. Alternatively, consider another scenario involving two people on a date, where one person suddenly gets up and leaves the table. There may be a key moment between event boundaries, such as a facial expression being a smile or a frown, that fundamentally captures the nature of that event and the way it will be interpreted and remembered. This moment may or may not be an event boundary.

In comparing key moments to event boundaries, paradigm-specific differences may account for which moments are categorized in terms of each phenomenon. That is, key moments are selected retrospectively, whereas event boundaries are typically judged in real-time. However, it is unlikely that these differences could fully account for our findings, for several reasons: First, both the segmentation and the key moment data were applied to a neutral fMRI-based dataset, where participants were only instructed to watch a movie and recall. Distinct neural signatures of event boundaries and key moments here suggest that there might be more than paradigm-specific differences between event boundaries and the key moments. Second, an exploratory univariate analysis investigating the role of event boundaries showed activation during event boundaries in the PMC and the hippocampus (Figure S3). The timing and profile of this univariate activation is highly consistent with prior work^5^, strongly suggesting that we did not simply fail to adequately capture event boundaries. Conversely, the activity for key moments peaked 6 seconds later, suggesting fundamental differences in activation profiles between key moments and event boundaries within the same task-relevant region of the brain. Nonetheless, future work could benefit from a systematic investigation of the paradigm-specific effects such as real-time versus retrospective frame selection.

Our approach provides a novel fine-grained method of comparing moment-to-moment activity patterns across people, as well as comparing them to recall activity patterns within participants. Prior work on episodic memory of movie stimuli has mostly looked at the average neural activity patterns of events during comprehension, and their relatedness to average recall patterns. This presents a challenge to understand the neural consequences at a more fine-grained level. Beyond yielding novel insights into event processing and recall within the present manuscript, we suggest that this approach can be used by future studies to address the importance of specific moments in establishing representations of extended events.

In general, we found sensitivity to key moments in posterior regions of the DMN. Given the roles of PMC^5,11,13,36^ and PHC^4,45–49^ in representing situations and contextual associations, it is sensible that these regions would selectively encode critical aspects of a narrative. Moreover, given that neural synchrony has been previously linked to shared understanding of stimuli ^7,11,36^, our findings suggest that key moments strongly capture context-sensitive information during movie viewing. In contrast, event boundaries resulted in decreased neural pattern synchrony among participants in these regions (Figure 4). One potential explanation for the desynchronization has to do with Event Segmentation Theory (EST). According to EST^50^, event boundaries arise when the prediction quality about the forthcoming experience is the lowest. It is possible that the prediction error may be experienced differently across participants, thus resulting in more idiosyncratic neural patterns at event boundaries. As a result, people differ from each other at event boundaries in the areas that track the content of the experience.

In contrast to PMC and PHC, mPFC and the hippocampus were more sensitive to event boundaries. Specifically, neural patterns were more synchronized across people at event boundaries in these regions, in contrast to the effects observed in the posterior regions of the DMN (Figure 4). Prior literature has suggested that the mPFC is involved in the forward projections of schemas at event boundaries^12,50^ and successful integration into episodic memory^21,51^. Similarly, the hippocampus is also shown to play a crucial role in segmenting continuous experiences and later retrieval^4,6,33,45,51–55^. Our results support and extend these findings, showing that these boundary preferences may exist above and beyond representing other components of an event. Finally, the participant’s activity patterns in the EVC and EAC regions also synchronized with each other at the key moments and event boundaries. It is sensible that we would observe this in sensory regions, given that participants all perceived the same content. Key moments may serve to heighten attention to meaningful pieces of information, which can sharpen perceptual processing ^56,57^ . We note, however, that effects in EVC and EAC did not extend to recall reinstatement. In contrast, event boundaries were not a significant predictor of the neural pattern synchrony in the EVC. One possible reason may be the locus of attentional focus. Prior work has shown that the participants’ eye movements are more dispersed at the event boundaries and eventually settle into the event’s progression^58^. As a result, the EVC may be processing different aspects of the scene at the event boundaries, thus resulting in a lack of synchronization of the neural activity around these time points.

During recall, neural patterns associated with key moments during encoding were overrepresented in neural activity in both the PMC and the mPFC. This was found in a moment-to-moment reinstatement analysis where the TRs with a high probability of being key moments were reinstated during recall, and also in a more conventional analysis where encoding patterns were simply weighted by key moment probabilities. In contrast, event boundary-weighted activity was not a significant predictor the recall activity. The interaction between key moments and event boundaries in both PHC and mPFC significantly predicted the recall activity, suggesting that both the event boundaries and key moments in these regions are crucial for event retrieval. This is consistent with the prior literature, wherein these two regions have been implicated in schema processing^12,50^ and contextual binding of information^4,45–49^, respectively. Findings in PHC and mPFC are also of note in the context of their lack of pattern reinstatement during event boundaries, as this is somewhat inconsistent with the idea that regions involved in episodic memory may selectively engage at boundaries^59,60^. Nonetheless, our results indicate that different brain regions important for encoding and retrieving event representations have unique dynamics as an event unfolds. Future work exploring functional connectivity analyses could help tease apart the distinct role of event boundaries and key moments in these regions and how they interact across various components of the Default Mode Network (DMN).

In sum, these findings provide novel evidence for key moments as important anchors of continuous events. People agree on which moments are critical for conveying the meaning of an extended narrative; these moments synchronize neural representations across individuals, and they dominate representational patterns that are present in the brain when people recall events.

### Behavioral study methods

#### The Storyboard paradigm

##### Participants

This study was administered in 4 parts. One hundred and ninety-eight participants across the United States were recruited online via Prolific (118 male, 77 female, 3 unknown; age 18-37, mean age = 29.2; approval rate between 90-100; standard sample) across the 4 parts of the study. Each part had a minimum of 48 participants and a maximum of 51 participants. Participants were provided with informed consent before the start of the study per the experimental procedures approved by the Danforth Institutional Review Board at Washington University in St. Louis. Each part lasted approximately 1 hour, and participants were compensated with $12 per hour prorated to their time. Given the novelty of the experiment design, no statistical methods were used to pre-determine sample sizes.

### Stimuli

The audio-visual content from the first 48 minutes of BBC’s *Sherlock S1E1: A Study in Pink* was used in this study. This stimulus has been used elsewhere in the literature^5,11,31^. The 48 minutes were broken down into 50 clips, each marked off at a coarse-grain event boundary as described in Chen et. al. 2017^11^ , and subsequently broken down into four parts. This was done to facilitate experiment loading for the participants on Prolific. Two of the clips in their stimuli were introductory cartoon segments that were used to facilitate fMRI data acquisition. These were excluded from our experiment, thus leaving us with 48 clips. Further, one more segment with introductory title cards was excluded. In addition, the first two segments at the beginning (the war scene in Afghanistan, and Watson subsequently waking up from a nightmare) were joined together to make one segment. Overall, this process resulted in 46 segments with the duration of each segment ranging from 11 to 185 seconds [SD = 41.6 seconds]. Finally, a 1-minute-long clip without audio from the movie *1917* was used as a practice example in the paradigm. The movie frame rate was 24 frames per second.

### Procedure

This experiment was administered online as a 4-part study. This was done to facilitate data loading and internet connectivity and to provide participants with enough time to create meaningful storyboards. Participants in each part were told they would watch a few clips from the BBC television crime drama *Sherlock*. After watching each clip, they used a custom-built interface to create the storyboards. The interface was programmed using the JsPsych 7.3.0 library^61^ and provided the participants with storyboard editing options such as Undo, Redo, Reorder, and Clear. Participants were instructed to recreate the underlying story in detail using the still images provided. A schematic of the experiment is illustrated in Figure 1. Participants were told they could select as many frames as they wanted to convey the underlying story in each clip. Upon completing each trial, participants were subsequently asked to rate on a Likert scale of 1 (low) – 7 (high) to determine their subjective ratings of the goodness of the generated storyboards. Finally, upon completing the experiment, they were also surveyed on Likert scales of 1 (low) – 5 (high) to determine whether they had any issues with video loading and had understood the instructions. A demonstration of the experiment can be found here (https://adibuoy23.github.io/event_representations/event_representation_paradigm.html).

### Event segmentation paradigm

The data for the event segmentation paradigm was obtained from another study^31^ that the same stimulus. A brief description of the materials for this study is noted below, however, more information can be found at this link: https://osf.io/preprints/psyarxiv/2dcq4.

### Participants

Thirty-nine participants were recruited from Cornell University for the event segmentation task involving the same Sherlock stimuli. The segmentation task was administered at both coarse and fine-grained with 21 participants (11 reported female and 10 reported Male; 18-39 years old; Mean = 20.95, SD = 4.40), and 18 participants (2 reported Female and 6 reported Male; 18-25 years old; Mean = 20.28, SD = 1.56) respectively.

### Stimulus

The first 48 minutes from BBC’s Sherlock S1:E1 A Study in Pink were also used to obtain event boundaries. The movie was divided into 10 clips overall, with the first 9 clips lasting 10 minutes each, and the 10^th^ clip lasting 3 minutes. The clips were cut at narrative shifts obtained from Chen et. al.^11^ Further, the ending of each clip was appended to the beginning of each clip to establish continuity during the task, and any repetitions in the content were subsequently excluded from the analysis. These appendices ranged from 9 to 17 seconds.

### Procedure

Participants in the coarse/fine segmentation condition were instructed to segment the activity into the largest or smallest units they found natural and meaningful. A shaping procedure consistent with prior literature on event segmentation^2,32,50^ was applied during the practice phase to both the coarse and fine-grained conditions. This was done primarily to ensure participants understood the task. A 91.1-second-long clip from the movie 3 Backyards was used as a practice clip for carrying out the shaping procedure.

### Data analysis

#### Data preprocessing for the storyboard paradigm

Participants who reported video loading issues or who did not understand the instructions (Likert scale responses <= 1) were excluded from further analysis. This operation excluded 9 participants across all 4 parts of the experiment, thus leaving 189 participants overall. On average, there were about 9.6 key moment frames per clip (S.D. = 3.9, min = 4, max = 19) across the 46 clips. To facilitate comparison between the key moments data and the event segmentation data, <u>t</u>imestamps in milliseconds were concatenated across all the clips to create a continuous time scale of the participant frame selection across the 48-minute movie.

### Data preprocessing for the event segmentation paradigm

Each participant’s segmentation time stamps were appended across the 10 clips to create a continuous timeline of the button presses for comparison with the key moments obtained from the storyboard paradigm. Segmentation data corresponding to the episode titles was removed to facilitate comparison with the storyboard data. This removed the segmentation data between the 2 min 51 sec and 3 min 16 sec mark. No other outlier removals based on button presses were implemented. This is consistent with recent methodological approaches to analyzing the event segmentation data ^62^. The continuous timeline was then divided into 46 clips to compare with the data from the storyboard paradigm. On average, there were 8.9 button presses in the fine-grained event boundary condition (S.D. = 5.9, min = 1, max = 25) – comparable with the key moment frames. There were 1.1 button presses in the coarse-grained event boundary condition (S.D = 0.3, min = 1, max = 3). Subsequent analyses comparing the key moments and event boundaries were restricted to the fine-grained condition, given the comparable statistics between the key moments and the fine-grained event boundaries.

### Comparing key moments and fine-grained event boundary distributions

For each participant, the time series of their key moments was convolved with a Gaussian kernel of Full Width Half Maximum (FWHM) ≈ 1.5 seconds to create a continuous distribution across all the clips. These individual density distributions were then averaged across the participants using a leave-one-out procedure to create a population density distribution, consistent with the prior literature analyzing segmentation data^32^. The generated individual and the leave-one-out population distributions were then split again into 46 clips for a linear mixed-effects analysis. Within each clip, the individual’s density distribution was correlated with the population density distribution to compute the key moment frame selection similarity among the participants. This resulted in a correlation score measuring the agreement between each participant and the rest of the population for each clip. This correlation will be referred to as the key moment-key moment correlation.

To test whether the agreement among participants was greater than expected by chance, a permutation analysis was conducted by picking key moment frames at random across the 48-minute movie. This was done by permuting the list of intervals between the two frames, and randomly sampling null frames between the intervals. This was done across the 48-minute video, and subsequently convolved with a Gaussian density kernel of FWHM ≈ 1.5 sec to generate a null distribution. This procedure was repeated 50 times for each subject, resulting in 50 null distributions for each subject. The continuous null distributions were also broken down into 46 clips, and each individual’s density distribution from the storyboard paradigm was then compared with the average of the 50 null distributions. This resulted in a distribution of null correlation scores for each participant and clip. This correlation score will be referred to as the key moment-null correlation.

Finally, the key moment distributions were compared to the fine-grained event boundary distributions by correlating individual storyboard density with the population event boundary density. This was done for every participant in the storyboard task and each clip. This operation yielded in a correlation score measuring the similarity between the storyboards and the event boundaries. This correlation score will be referred to as the key moment-event boundary correlation for the remainder of the document.

Similar operations were implemented for event boundaries where individual densities were compared with population event boundary density, the simulated null distribution density, and the key moment population density distributions. These correlations will be referred to as event-boundary-event-boundary, event-boundary-null, and event-boundary-key moment correlations for the remainder of the document. A schematic illustrating the above analyses is shown in Figure 2A. See Supplementary Information for more information on the data analysis and an extended version of the results.

### fMRI methods

#### Participants & stimuli

We used a publicly available fMRI dataset^11^ from 17 participants who watched the first 48 minutes of Sherlock S1:E1 (“A Study in Pink”) while undergoing scanning (https://openneuro.org/datasets/ds001132/versions/00003). After viewing, participants completed a free recall task, recounting the movie in as much detail as possible. On average, participants recalled approximately 34.4 clips (SD = 6) with a mean duration of 21.7 minutes (SD = 8.9). The obtained transcripts were used to annotate the neural recall data and map it to the specific clips.

The MRI data were collected on a 3T full-body scanner (Siemens Skyra) with a 20-channel head coil. Functional images were acquired using T2*-weighted an echo planar imaging (EPI) pulse sequence (TR = 1.5 sec, TE = 28 msec, flip angle 64) with whole-brain coverage (27 slices of 4 mm thickness, 3 x 3 mm^2^ in-plane resolution, 192 x 192 mm^2^ FOV, ascending interleaved acquisition). High resolution anatomical images were acquired using a T1-weighted MPRAGE pulse sequence (0.89 mm^3^ resolution).

### Data Analysis

#### Preprocessing

The fMRI data were preprocessed using FSL [http://fsl.fmrib.ox.ac.uk/fsl] following the procedure described by Chen et al^11^. Preprocessing included slice timing correction, motion correction, linear detrending, high-pass filtering (140s cutoff), spatial normalization to MNI space, and resampling to 3mm isotropic voxels. A 6 mm spatial smoothing kernel was applied, and voxel time series were z-sored to enable inter-subject comparisons. To account for hemodynamic lag, all data were shifted by 3 TRs (4.5s). Motion was minimized using head padding, and one subject’s scan, which had missing data near the end, was padded to match the others in length.

The data from the two cartoon segments in this dataset were subsequently excluded from the analysis. In addition, the neural data pertaining to the encoding and recall of the title card clip were also excluded from subsequent fMRI analyses. The remainder encoding data was re-aligned with the timestamps of the 46 clips obtained from the storyboard paradigm, and any recall data present for these 46 clips was considered in our subsequent analyses.”

### Defining the Regions of Interest (ROIs)

The ROIs that were used in this analysis include the DMN regions, namely the PMC, PHC, PRC, and mPFC. These regions were selected because prior studies implicated a distinct role for them in naturalistic movie comprehension^5,7,11,12,33^. These ROIs were obtained from Chen et al. 2017.

### Comparing the key moment and event boundary distributions with the neural data

Each fMRI participant (n=17) had neural data that spanned 1976 TRs during the movie viewing. The TRs corresponding to the title cards from this data were excluded from analysis <u>because</u> key moments and event boundaries were not collected <u>for</u> these clips. For each TR, the probability of identification as a key moment was calculated by integrating over the estimated probability density of identification as a key moment for the 1.5-s length of the TR. (See *Comparing key moments and fine-grained event boundary distributions*, above.) The same thing was done to calculate the probability of being an event boundary.

Next, a constant 0.001 value was added to the key moment and event boundary probabilities to have non-zero values for the log transformation. Finally, the log-transformed probabilities were z-scored across all the TRs to address skewness in the two distributions and to facilitate subsequent neural analyses comparing ISPS and the generated probability distributions. See Supplementary Information for more information on the data analysis and an extended version of the results.

## Supplementary Information

Supplementary analysis S1: Permutation analysis comparing shuffled vs intact distributions of key moments and event boundaries in predicting the ISPS:

We tested whether key moments (KMs) and event boundaries (EBs) reliably predicted ISPS activity by comparing intact probability distributions against shuffled nulls. For each movie clip (segmented by experimenter-identified coarse boundaries), we generated 1000 null distributions using a phase-scrambling procedure that preserved interval durations. Each intact and shuffled distribution was then entered as a fixed effect in linear mixed-effects models (lmerTest in R) predicting ISPS, with participant and clip as random intercepts. The resulting beta weights from the intact models were compared against the shuffled null distribution.

Intact KM betas exceeded shuffled nulls across most ROIs (PMC: Δ = 0.007, 95% CI [0.001, 0.01], p = .0009; PHC: Δ = 0.006, [0.002, 0.01], p = .0009; MPFC: Δ = 0.0037, [-0.001, 0.008], p = .051; Hippocampus: Δ = 0.0043, [0.0001, 0.008], p = .011; PRC: Δ = -0.001, [-0.009, 0.0001], p = .018; EVC: Δ = 0.01, [0.005, 0.016], p = .0009; EAC: Δ = 0.015, [0.01, 0.02], p = .0009). In contrast, EB betas were generally lower than shuffled nulls, with the exception of MPFC and hippocampus, where intact EBs were higher (PMC: Δ = -0.01, [-0.016, -0.005], p = .0009; PHC: Δ = -0.005, [-0.01, -0.006], p = .01; MPFC: Δ = 0.007, [0.002, 0.011], p = .0049; Hippocampus: Δ = 0.007, [0.003, 0.011], p = .0009; PRC: Δ = -0.003, [-0.008, 0.02], p = .12; EAC: Δ = 0.015, [0.01, 0.02], p = .0009; EVC: Δ = -0.004, [-0.01, 0.002], p = .086). These results confirm and extend the main ISPS findings: intact KMs and EBs carry distinct neural signatures during movie viewing (see Figure S 1) for more details.

**Figure S1:**
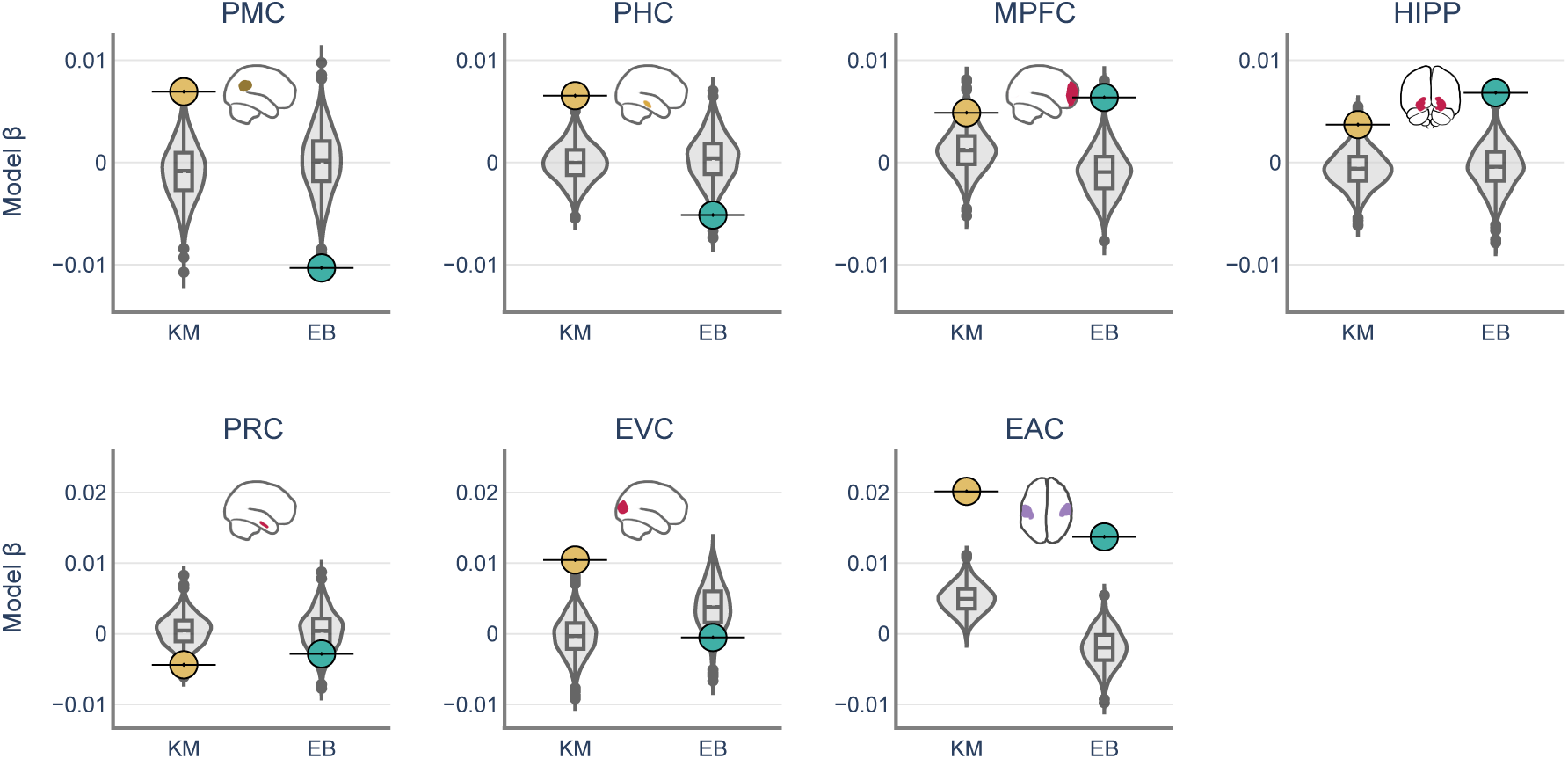
Comparing the permuted distributions of key moments and event boundaries to their intact counter parts. Individual colored dots (yellow: key moments (KM), green: event boundaries(EB)) indicate the beta value of the intact probability distribution in predicting the ISPS, whereas the gray distributions indicate the beta values of the randomized probability distributions in predicting the ISPS signal.

Supplementary analysis S2: Key moment weighted movie-viewing encoding neural activity is over-represented in the average neural recall activity:

We tested whether neural activity at key moments (KMs) during encoding was preferentially reinstated during recall (Figure S 2 A). For each TR, encoding patterns were weighted by the probability of a KM or event boundary (EB), and these weighted averages were compared with recall activity in a linear mixed-effects model, with subjects and clips modeled as random effects.

KM-weighted encoding significantly predicted recall in PMC (β = 0.07, p = .004) and mPFC (β = 0.06, p = .047), whereas EB-weighted encoding did not predict recall in any ROI (Figure S 2B). The KM × EB interaction was significant in mPFC (β = –0.018, p < .001) and PHC (β = –0.005, p < .001; Table 5), suggesting a downregulation of reinstatement at event boundaries, even when those boundaries coincided with KMs.

**Table 5:**
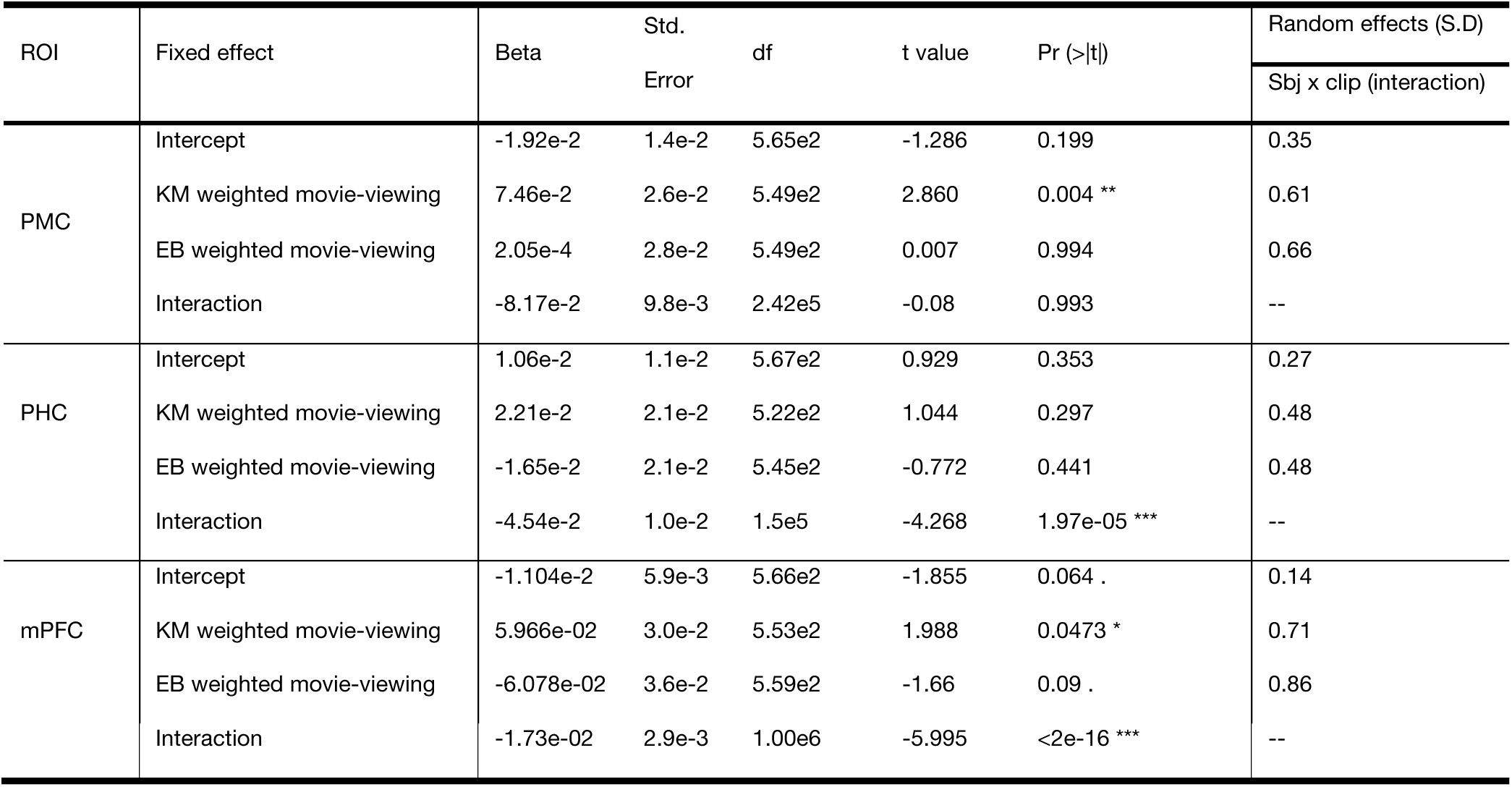
Linear mixed-effects models evaluating the recall activity as a function of the key moment and event boundary weighted movie-viewing activity. Three ROIs Posterior Medial Cortex (PMC), Parahippocampal cortex (PHC), and the medial Prefrontal Cortex (MPFC) were used this analysis.

In sum, KMs during movie viewing predicted recall activity above and beyond EBs, underscoring their role in shaping the neural representations that support memory.

**Figure S2:**
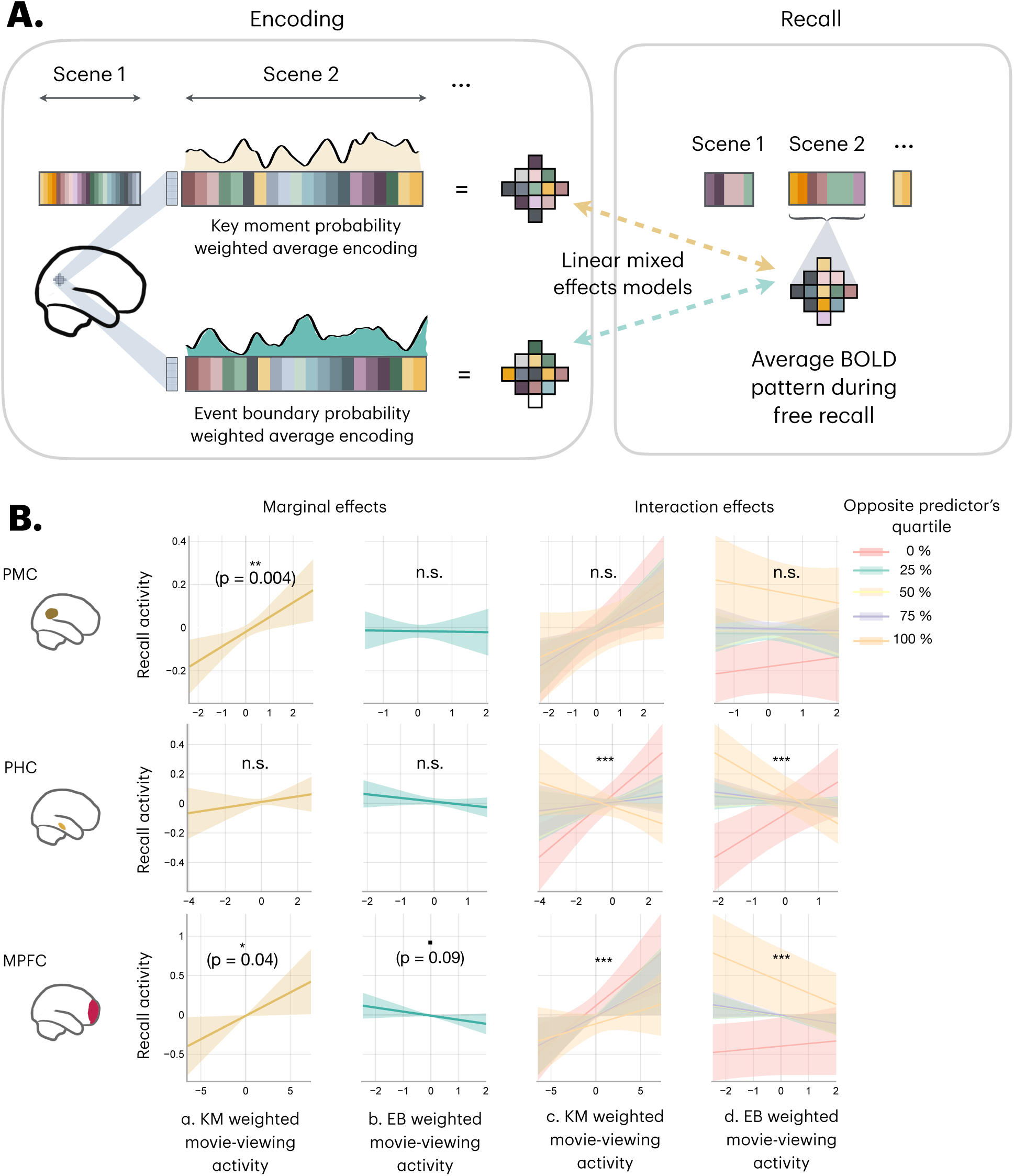
Top: Schematic comparing the neural activation of recall to the movie viewing data. To check for the effect of key moments on recall, the movie-viewing neural data has been weighted by both key moment and event boundary probabilities and compared with the average neural pattern of activity during free recall of each scene, provided the scene was recalled by a given participant. Bottom: Linear mixed-effects models comparing key-moment (KM) and event boundary (EB) weighted movie-viewing data against the recall activity patterns. Each row corresponds to an ROI. Column a: Recall activity as a function of key moment weighted movie-viewing activity. KM-specific movie-viewing activity significantly predicted the recall activity in PMC and MPFC. Column b: Recall activity as a function of EB-weighted movie-viewing activity. EB-specific movie-viewing activity was marginally significant in predicting the recall activity in MPFC. Column c: Recall activity as a function of KM-weighted movie-viewing activity grouped by different quartiles of predicted EB probabilities using LMER model. Column d: Recall activity as a function of EB-weighted movie-viewing activity grouped by different quartiles of predicted KM probabilities using LMER model. There were significant interactions between KM and EB weighted movie-viewing data in predicting the recall data in PHC, and MPFC. The solid lines indicate the estimates of the linear mixed effects model, and the shaded regions indicate the 95% CI of the estimates.

Supplementary analysis S3: Overall, BOLD activity in the Posterior Medial Cortex (PMC), Medial Prefrontal Cortex (MPFC), and the Hippocampus is higher at Event Boundaries:

We examined overall BOLD activity in PMC, mPFC, and hippocampus using a univariate FIR analysis (see Methods). Voxel wise movie-viewing activity was z-scored within subjects, and predictors were defined by peaks in the key moment (KM) and event boundary (EB) probability distributions. FIR models were fit to the z-scored data, shifted 3 TRs for hemodynamic lag, and linear mixed-effects models tested lag effects with subjects and voxels as random effects.

Both KMs and EBs showed significant main effects of lag in PMC (EB: χ²(13) = 300.08, p < .0001; KM: χ²(13) = 327.05, p < .0001), mPFC (EB: χ²(13) = 130.58, p < .0001; KM: χ²(13) = 62.97, p < .0001), and hippocampus (EB: χ²(13) = 228.68, p < .0001; KM: χ²(13) = 67.00, p < .0001) (Figure S 3). EB-related activity consistently peaked around lag 0, while KM-related peaks varied across regions (0–6s).

These results replicate prior reports of boundary-evoked BOLD activity increases^4,5^ and show that KMs also elicit region-specific univariate responses.

**Figure S3:**
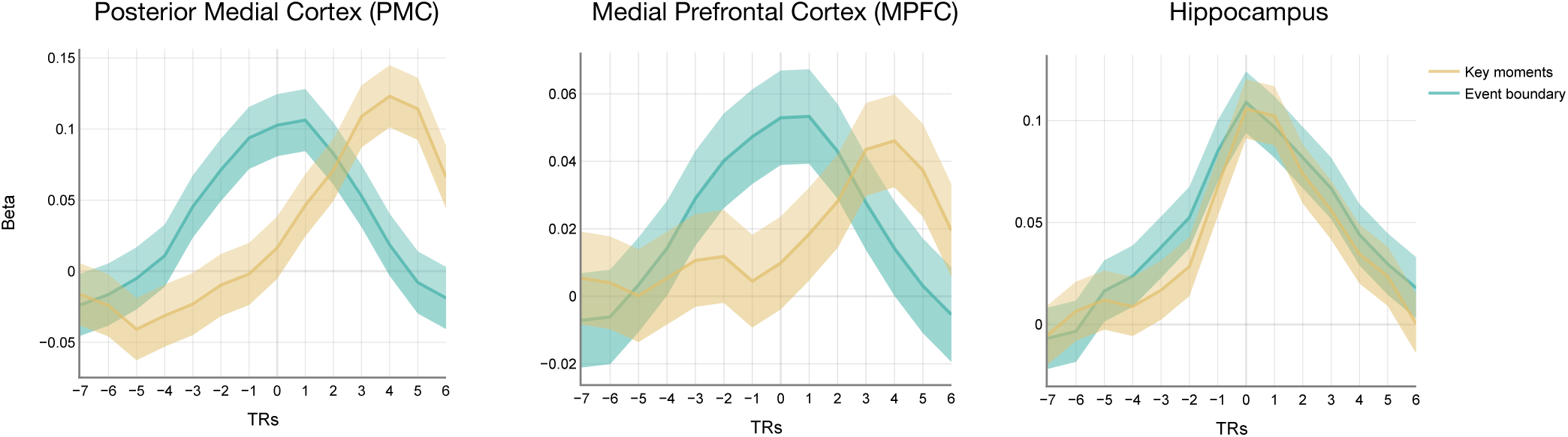
Univariate analysis comparing the BOLD activity around event boundaries and key moments in the Posterior Medial Cortex (PMC), Medial Prefrontal Cortex (MPFC), and the Hippocampus (Hipp). Beta values represent the degree of activation across the voxel relative to the baseline and are obtained by the Finite Impulse Response (FIR) model. The solid line represents the linear mixed effects model estimates of beta values. The shaded lines represent the 95% CI.

Supplementary analysis S4: The effect of unique key moment and event boundary instants in predicting the ISPS:

We checked whether the unique probability of being a KM or EB predicted the ISPS. The analysis for this followed the same procedures as the ISPS analysis (Figure 3). Except, the predictor variables were the unique KM and unique EB probabilities. The unique probabilities of both the metrics were obtained by regressing out the common variance between the two distributions. A schematic of the analysis is displayed below (Figure S 4). The results replicated the findings of the ISPS analysis (Figure S 5, Table S 1).

**Figure S4:**
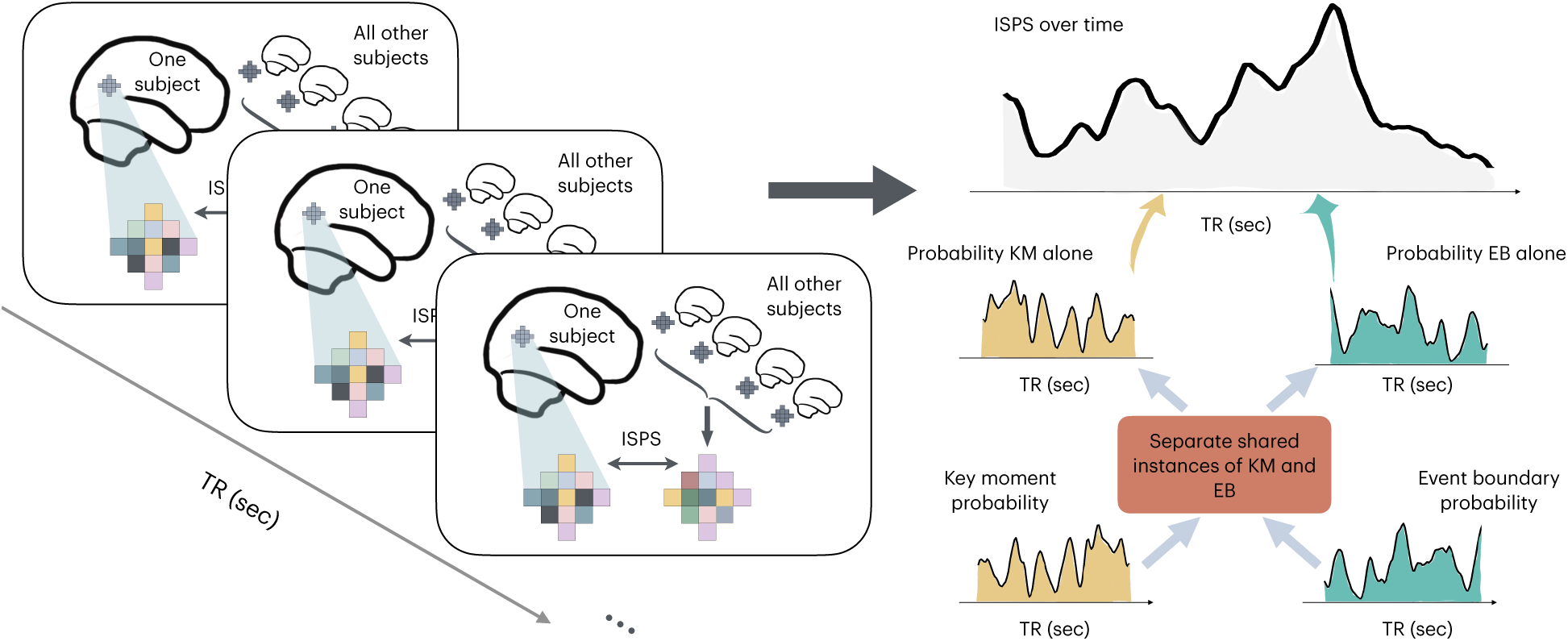
Intersubject Neural Pattern Similarity schematic as a function of unique KM and unique EB probability distributions. The unique KM and EB probability distributions were obtained by regressing out the shared components between the two, and the left-over residual from the linear model is used to obtain the unique components of the KM and EB distributions, respectively.

**Figure S5:**
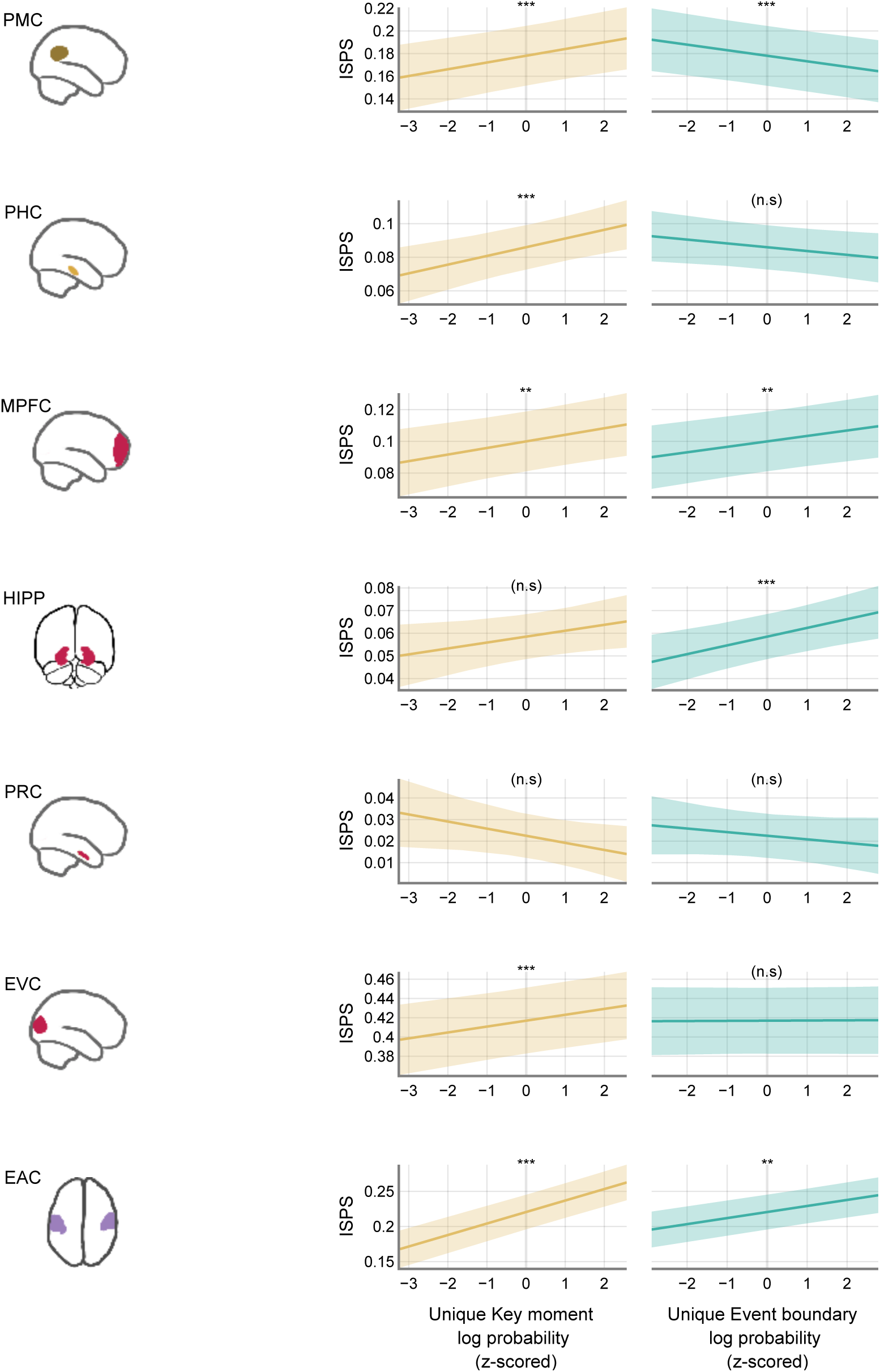
Comparing Inter-Subject neural Pattern Similarity (ISPS) against the unique key moment and event boundary probabilities. The first column indicates the ROI that was examined. The second column plots the relationship between the ISPS and the unique key moment probability distribution after log-transformed and z-scored. The third column shows the relationship between ISPS and the unique event boundary probability.

**Table S1:**
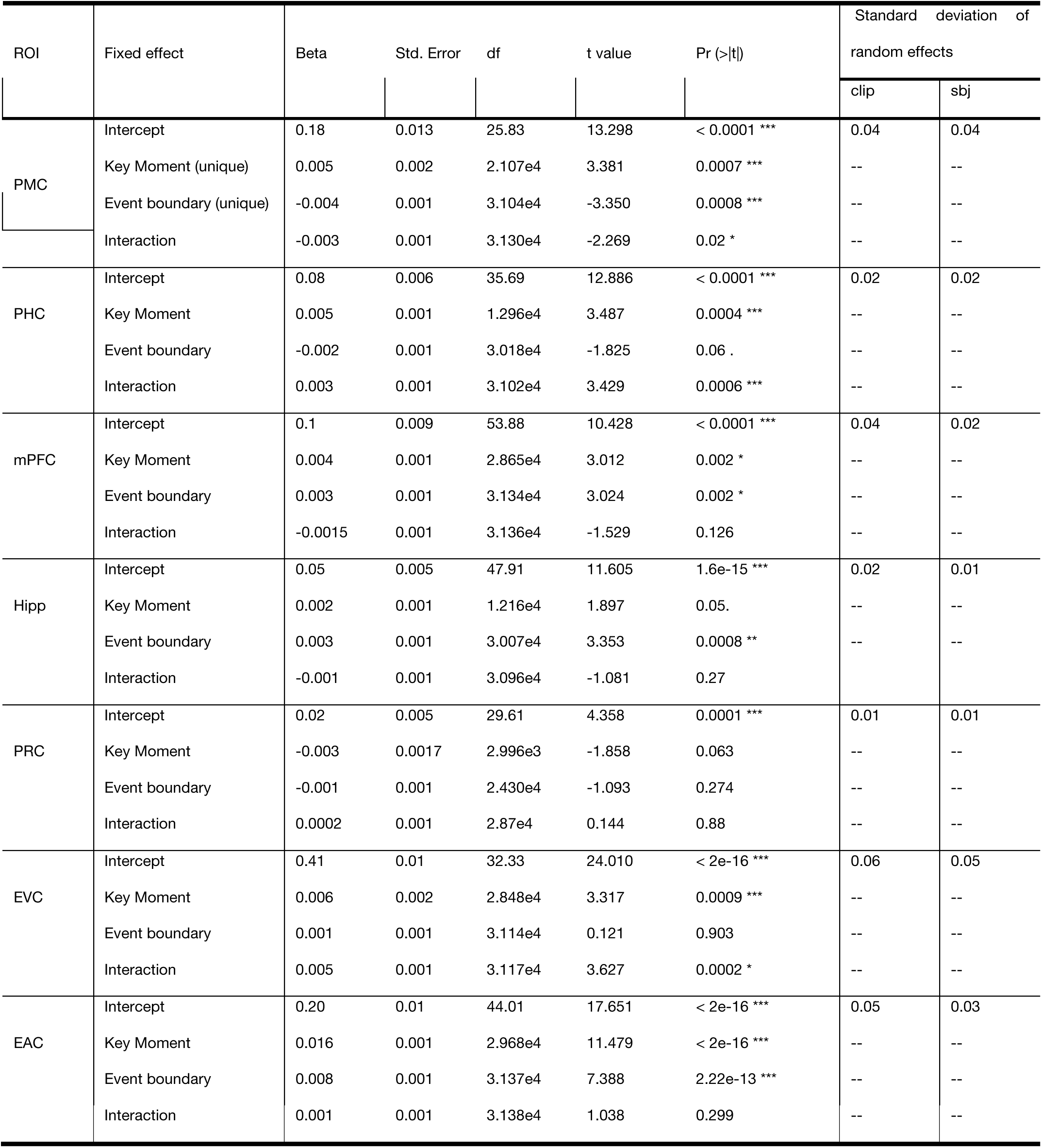
Linear mixed-effects models comparing the ISPS activity over time with unique key moment and event boundary probability distributions.

### Behavioral results (extended)

Participants agree on key moments within continuous events, which are partially overlapping with but dissociable from event boundaries

A. *Participants agreed on the locations of key moments*: A linear mixed effects model analysis using Restricted Maximum Likelihood (REML) compared the KM-KM correlations against the KM-null correlations. The correlations were the dependent variables, and the condition (key moment vs. null) was the fixed effect. The contrasts for condition were dummy coded. Participants and clips were modeled as random effects with a maximal random effects structure of random intercepts and random slopes of condition. This analysis revealed a significant main effect of the key moment-key moment condition (β = 0.40, SEM = 0.03, t(53.5) = 12.627, p < 0.001, Figure 2B). Random intercepts of participants accounted for 0.4% of the total variance (SD = .06); random slopes for the condition accounted for 1.5% of the total variance (SD = 0.12). Random intercepts for clips accounted for 4% of the total variance (SD = 0.21); random slopes for the condition accounted for 4% of the total variance. (S.D = 0.2). This analysis suggests that participant frame selections greatly agree with each other in the storyboard task. The distribution of the correlations for the KM-KM and KM-null conditions can be seen in Figure 2B.

B. *Participants agreed on the locations of event boundaries.* A linear mixed effects model analysis using REML compared the EB-EB correlations against the EB-null correlations. The correlations were the dependent variables, and the condition (event boundary vs. null) was the fixed effect. The contrasts for condition were dummy coded. Participants and clips were modeled as random intercepts and random slope of the condition was also included, similar to the KM analysis above. This analysis also revealed a significant main effect of the EB-EB condition (β = 0.25, SE = 0.03, *t*(26.72) = 7.809, *p* < 0.001). Random intercepts of participants accounted for 1% of the total variance (SD = .10); random slopes for the condition accounted for 1% of the total variance (SD = 0.10). Random intercepts for clips accounted for 1% of the total variance (SD = 0.11); random slopes for the condition accounted for 1% of the total variance (S.D = 0.11). These results suggest that participant button press timestamps were similar to each other compared to the random null distribution, thus replicating the existing findings on event segmentation^32,63^.

*C. The locations of key moments differed significantly from those of event boundaries.* A linear mixed effects model analysis using REML compared the key moment-key moment correlations to the key moment-event boundary correlations. The correlations were the dependent variables, and the condition (KM-KM vs. KM-EB) was the fixed effect. The contrasts for condition were dummy coded. Participants and clips were modeled as random effects with clips including random intercepts and the condition as a random slope. Random effects of participants only included the random intercept. This was the maximal random effects structure that led to convergence without a singular fit. This analysis revealed a significant main effect of the key moment-key moment condition (β = 0.25, S.E = 0.01, *t*(45.20) = 13.361, *p* < 0.001). Random intercepts of participants accounted for 0.6% of the total variance (SD = .08. Random intercepts for clips accounted for 2% of the total variance (SD = 0.14); random slopes for the condition accounted for 1% of the total variance (S.D = 0.12). Similarly, the linear mixed effects analysis comparing event boundary–event boundary correlations against the EB-KM correlations also revealed a significant main effect of the EB-EB condition (β = 0.05, SE = 0.01, *t*(44.45) = 3.632, *p* = 0.0007). The random effects structure in this analysis involved participants and clips as random intercepts. The condition was modeled as a random slope for the clips condition. This was the maximal random effects structure that led to convergence without a singular fit. Random intercepts of participants accounted for 3% of the total variance (SD = .06. Random intercepts for clips accounted for 1% of the total variance (SD = 0.10); random slopes for the condition accounted for 0.3% of the total variance (S.D = 0.06). The distribution plots comparing KM-KM correlations to EB-KM correlations and vice versa are also shown in Figure 2. Together, these results revealed that the key moment frames selected by the participants were highly similar to each other, and significantly differed from the null frames and the fine-grained event-boundaries.

#### fMRI results (extended)

##### Movie viewing

###### Neural activity patterns synchronize across people at key moments

We computed the ISPS to compare how similar the BOLD activity patterns are across participants over time. This was done by correlating the activity pattern for a given subject and TR in a clip with the average activity pattern from the other 16 subjects (n=17) for the same TR and clip. This procedure was repeated for all 46 clips. The ROIs used in this analysis largely focused on the regions within the DMN. These regions included the PMC, PHC, mPFC, hippocampus, and PRC^7,12,12,31,33,35,38,53^. In addition, EVC and EAC were also used as control regions in the analysis. A schematic of this analysis is described in Figure 3.

*A. Inter-subject neural pattern synchrony reliably tracked the content of the movie during encoding:* The ISPS derived from the brain activity patterns was analyzed to see if it reflects the contents of the underlying movie viewing experience. To do this, a null correlation metric similar to the ISPS was obtained. This was done by measuring each subject’s brain activity pattern in a given TR and clip and comparing it to the rest of the subjects’ average brain activity pattern for a TR of the same index for a flipped clip. For example, a given subject’s activity pattern in TR “1” would be correlated to the rest of the subjects’ average activity pattern in the TR “N-1”, where N is the total number of TRs in the given clip. The two correlation distributions, i.e., the ISPS and the null distribution, were then compared to see if they were significantly different from each other. The rationale behind this analysis is that if the brain activity patterns are accurately tracking the content of the movie, they would be significantly different from the null condition. A linear mixed effects model with the correlation as the dependent variable and the correlation type (ISPS vs. null) as the fixed effects was used to compare for statistical differences between the two metrics. Subjects and clips were modeled as random effects with the correlation type variable as the random slope. The best model was determined for each ROI by tracking model convergence and singularity fit. This analysis revealed a significant difference between the ISPS and the null correlation type across all the seven ROIs described above. These results suggest that the ISPS signal across the participants captured reliable information about the movie viewing experience and was not due to structural artifacts in the brain. The output of the linear mixed effects model for the 7 ROIs is summarized in Table 3, and the results can be seen in Figure 4B (middle column).

*B. Comparing inter-subject neural pattern synchrony over time against key moment and event boundary probability distributions*: To analyze whether the neural patterns were synchronized around the key moments vs. event boundaries, the ISPS signal for each subject and clip was compared to the key moment and event boundary probability distributions (log transformed and z-scored) within that clip; see fMRI Methods for more information on aligning the granularity of both key moment and event boundary distributions to the neural data. An LME model was implemented with the ISPS as the dependent variable and the key moment and event boundary probabilities as the predictor variables. The 17 fMRI participants and the 46 clips were modeled as the random effects. The interaction terms in the fixed effects and the inclusion of the random slopes into the models were determined by the model selection procedures following model convergence and singularity warnings. We note that this analysis could be susceptible to inflation of effects due to temporal autocorrelation in the large number of time points. We also report another bootstrapping analysis (see Supplemental information S1) to empirically determine the significance values. Overall, the results showed a significant positive correlation between the key moment probabilities and the ISPS scores in the PMC and PHC. However, event boundaries had a significant negative effect on the ISPS in these regions. See Figure 4B for more information and Table 2 for the results of the linear mixed effects models across various ROIs.

### Free recall

#### Key moments dominate neural pattern reinstatement during event recall

##### Neural pattern reinstatement analysis

To investigate the contribution of the key moments in the recall, we investigated whether their corresponding neural patterns during encoding were reinstated during recall. We correlated the neural pattern for each TR during movie viewing with the average recall neural pattern over time. In other words, this analysis measured how much of the pattern corresponding to each TR was reinstated during the overall recall activity. The computed signal is termed as neural pattern reinstatement (NPR) over time.

*A. Neural pattern reinstatement measures reliable signals for each clip across participants:* To measure whether the NPR signal over time was a reliable signal of interest, we compared the NPR signal over time against the null distribution. This is referred as within-clip condition in Figure 5. The null distribution of the NPR was determined by taking the neural pattern for each clip and TR in an ROI during movie encoding and comparing it with the average recall pattern for a different clip from the same ROI during recall. This is referred as across-clips in Figure 5. The null NPR should therefore reflect stable spatial variance in the ROI throughout movie viewing, but no scene-specific patterns. A linear mixed effects model with the correlation as the dependent variable and the correlation type (within-clip vs. across-clips) as the fixed effects was used to compare for statistical differences between the two metrics. Subjects and clips were modeled as random intercepts, and correlation type was used as random slope. The best model was determined for each ROI by tracking model convergence and singularity fit. This analysis revealed a significant difference between the NPR and the null correlation type in the PMC, mPFC, and PHC (see Table 3 and Figure 5). No significant difference between the two distributions was observed for the other ROIs. These results suggest that the NPR signal across the participants reflected reliable reinstatement of moment-to-moment neural patterns at the time of encoding during recall.

*B. Comparing the neural pattern reinstatement signal with key moment and event boundary probabilities.* To measure whether the key moment movie viewing patterns were reliably reinstated during recall, a linear mixed effects model was implemented with the NPR as the dependent variable. The log-transformed and z-scored key moment and event boundary probabilities and their interactions were modeled as independent variables. The 17 fMRI participants and the 46 clips were modeled as random effects. The 46 clips were nested within each subject. This was done for two reasons: 1. Each participant’s recall of the scene could be unique. 2: Not every participant recalled all 46 clips. Further, the inclusion of the random slopes into the models was determined by the model selection procedures following model convergence and singularity warnings. The results showed a significant positive effect of the key moment probabilities in predicting the NPR scores within the PMC. There was no significant main effect of event boundaries within this ROI. Further, the interaction was also not a significant predictor of the NPR signal within this region. The same analysis was repeated for both PHC, and mPFC regions. None of the predictor variables were significant predictors in both the ROIs (see Table 4; Figure 6).

