## Supplemental Information for "Key moments in naturalistic events synchronize neural activity patterns and dominate memory reinstatement"

Supplementary Information:

<u>Supplementary analysis S1: Permutation analysis comparing shuffled vs intact distributions of key moments and event boundaries in predicting the ISPS:</u>

We tested whether key moments (KMs) and event boundaries (EBs) reliably predicted ISPS activity by comparing intact probability distributions against shuffled nulls. For each movie clip (segmented by experimenter-identified coarse boundaries), we generated 1000 null distributions using a phase-scrambling procedure that preserved interval durations. Each intact and shuffled distribution was then entered as a fixed effect in linear mixed-effects models (lmerTest in R) predicting ISPS, with participant and clip as random intercepts. The resulting beta weights from the intact models were compared against the shuffled null distribution.

Intact KM betas exceeded shuffled nulls across most ROIs (PMC: Δ = 0.007, 95% CI [0.001, 0.01], p = .0009; PHC: Δ = 0.006, [0.002, 0.01], p = .0009; MPFC: Δ = 0.0037, [-0.001, 0.008], p = .051; Hippocampus: Δ = 0.0043, [0.0001, 0.008], p = .011; PRC: Δ = -0.001, [-0.009, 0.0001], p = .018; EVC: Δ = 0.01, [0.005, 0.016], p = .0009; EAC: Δ = 0.015, [0.01, 0.02], p = .0009). In contrast, EB betas were generally lower than shuffled nulls, with the exception of MPFC and hippocampus, where intact EBs were higher (PMC: Δ = -0.01, [-0.016, -0.005], p = .0009; PHC: Δ = -0.005, [-0.01, -0.006], p = .01; MPFC: Δ = 0.007, [0.002, 0.011], p = .0049; Hippocampus: Δ = 0.007, [0.003, 0.011], p = .0009; PRC: Δ = -0.003, [-0.008, 0.02], p = .12; EAC: Δ = 0.015, [0.01, 0.02], p = .0009;

EVC: Δ = -0.004, [-0.01, 0.002], p = .086). These results confirm and extend the main ISPS findings: intact KMs and EBs carry distinct neural signatures during movie viewing (see Figure S 1) for more details.

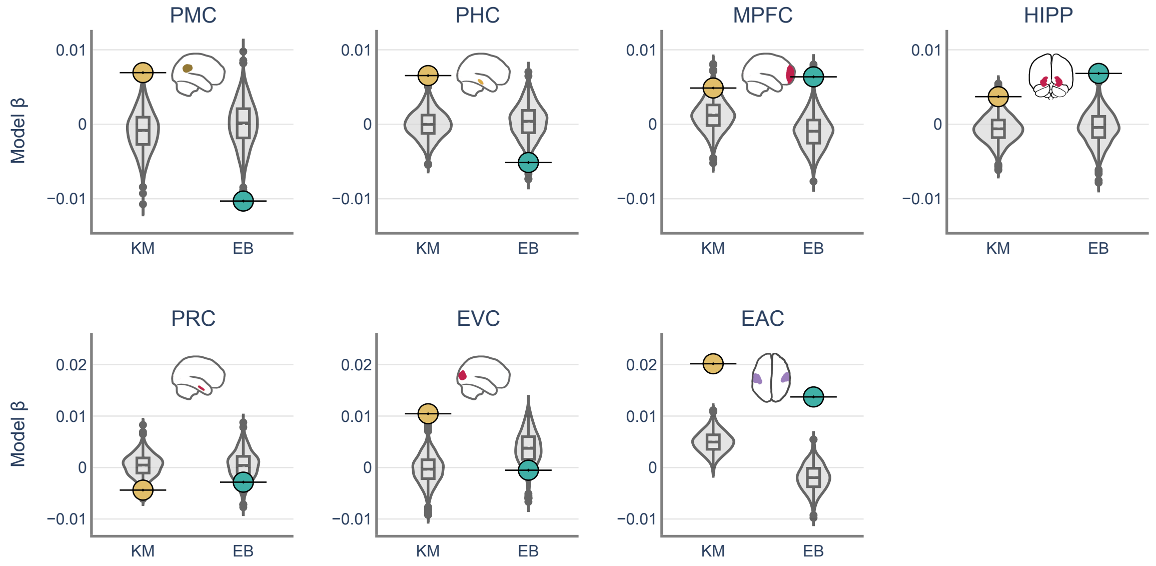


*Figure S 1: Comparing the permuted distributions of key moments and event boundaries to their intact counter parts. Individual colored dots (yellow: key moments (KM), green: event boundaries(EB)) indicate the beta value of the intact probability distribution in predicting the ISPS, whereas the gray distributions indicate the beta values of the randomized probability distributions in predicting the ISPS signal.*

<u>Supplementary analysis S2: Key moment weighted movie-viewing encoding neural activity is over-represented in the average neural recall activity</u>:

We tested whether neural activity at key moments (KMs) during encoding was preferentially reinstated during recall (Figure S 2 A). For each TR, encoding patterns were weighted by the probability of a KM or event boundary (EB), and these weighted averages were compared with recall activity in a linear mixed-effects model, with subjects and clips modeled as random effects.

KM-weighted encoding significantly predicted recall in PMC ($\beta = 0.07$, $p = .004$) and mPFC ($\beta = 0.06$, $p = .047$), whereas EB-weighted encoding did not predict recall in any ROI (Figure S 2B). The KM × EB interaction was significant in mPFC ($\beta = -0.018$, $p < .001$) and PHC ($\beta = -0.005$, $p < .001$; Table 5), suggesting a downregulation of reinstatement at event boundaries, even when those boundaries coincided with KMs.

In sum, KMs during movie viewing predicted recall activity above and beyond EBs, underscoring their role in shaping the neural representations that support memory.

*Table 5: Linear mixed-effects models evaluating the recall activity as a function of the key moment and event boundary weighted movie-viewing activity. Three ROIs Posterior Medial Cortex (PMC), Parahippocampal cortex (PHC), and the medial Prefrontal Cortex (MPFC) were used this analysis.*

| ROI | Fixed effect | Beta | Std. Error | df | t value | Pr (>\|t\|) | Random effects (S.D) |
|---|---|---|---|---|---|---|---|
| | | | | | | | Sbj x clip (interaction) |
| PMC | Intercept | -1.92e-2 | 1.4e-2 | 5.65e2 | -1.286 | 0.199 | 0.35 |
| | KM weighted movie-viewing | 7.46e-2 | 2.6e-2 | 5.49e2 | 2.860 | 0.004 ** | 0.61 |
| | EB weighted movie-viewing | 2.05e-4 | 2.8e-2 | 5.49e2 | 0.007 | 0.994 | 0.66 |
| | Interaction | -8.17e-2 | 9.8e-3 | 2.42e5 | -0.08 | 0.993 | -- |
| PHC | Intercept | 1.06e-2 | 1.1e-2 | 5.67e2 | 0.929 | 0.353 | 0.27 |
| | KM weighted movie-viewing | 2.21e-2 | 2.1e-2 | 5.22e2 | 1.044 | 0.297 | 0.48 |
| | EB weighted movie-viewing | -1.65e-2 | 2.1e-2 | 5.45e2 | -0.772 | 0.441 | 0.48 |
| | Interaction | -4.54e-2 | 1.0e-2 | 1.5e5 | -4.268 | 1.97e-05 *** | -- |
| mPFC | Intercept | -1.104e-2 | 5.9e-3 | 5.66e2 | -1.855 | 0.064 . | 0.14 |
| | KM weighted movie-viewing | 5.966e-02 | 3.0e-2 | 5.53e2 | 1.988 | 0.0473 * | 0.71 |
| | EB weighted movie-viewing | -6.078e-02 | 3.6e-2 | 5.59e2 | -1.66 | 0.09 . | 0.86 |
| | Interaction | -1.73e-02 | 2.9e-3 | 1.00e6 | -5.995 | <2e-16 *** | -- |

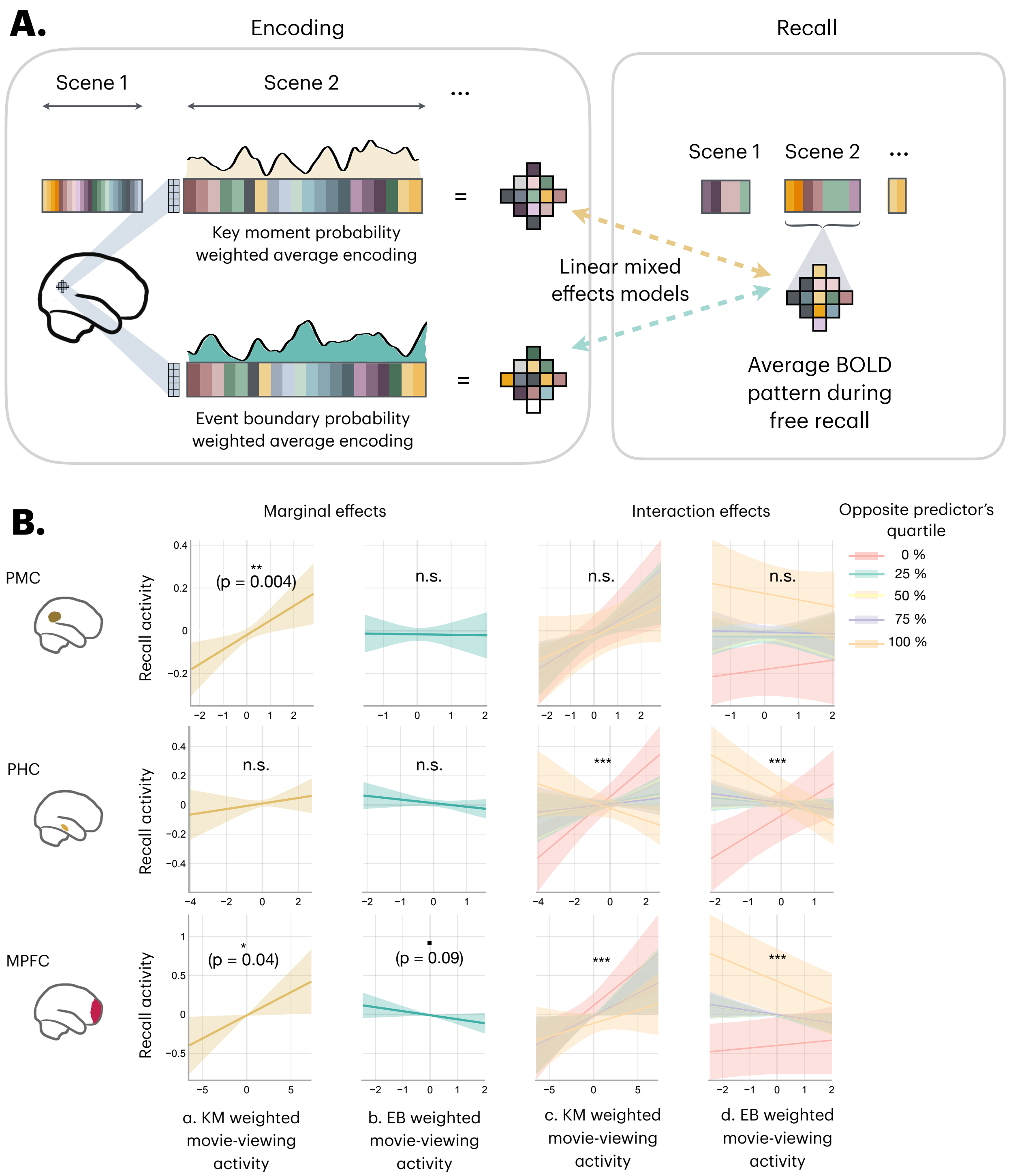


*Figure S 2: Top: Schematic comparing the neural activation of recall to the movie viewing data. To check for the effect of key moments on recall, the movie-viewing neural data has been weighted by both key moment and event boundary probabilities and compared with the average neural pattern of activity during free recall of each scene, provided the scene was recalled by a given participant. Bottom: Linear mixed-effects models comparing key-moment (KM) and event boundary (EB) weighted movie-viewing data against the recall activity patterns. Each row corresponds to an ROI. Column a: Recall activity as a function of key moment weighted movie-viewing activity. KM-specific movie-viewing activity significantly predicted the recall activity in PMC and MPFC. Column b: Recall activity as a function of EB-weighted movie-viewing activity. EB-specific movie-viewing activity was marginally significant in predicting the recall activity in MPFC. Column c: Recall activity as a function of KM-weighted movie-viewing activity grouped by different*

*quartiles of predicted EB probabilities using LMER model. Column d: Recall activity as a function of EB-weighted movie-viewing activity grouped by different quartiles of predicted KM probabilities using LMER model. There were significant interactions between KM and EB weighted movie-viewing data in predicting the recall data in PHC, and MPFC. The solid lines indicate the estimates of the linear mixed effects model, and the shaded regions indicate the 95% CI of the estimates.*

<u>Supplementary analysis S3: Overall, BOLD activity in the Posterior Medial Cortex (PMC), Medial Prefrontal Cortex (MPFC), and the Hippocampus is higher at Event Boundaries</u>:

We examined overall BOLD activity in PMC, mPFC, and hippocampus using a univariate FIR analysis (see Methods). Voxel wise movie-viewing activity was z-scored within subjects, and predictors were defined by peaks in the key moment (KM) and event boundary (EB) probability distributions. FIR models were fit to the z-scored data, shifted 3 TRs for hemodynamic lag, and linear mixed-effects models tested lag effects with subjects and voxels as random effects.

Both KMs and EBs showed significant main effects of lag in PMC (EB: $\chi^2(13) = 300.08$, $p < .0001$; KM: $\chi^2(13) = 327.05$, $p < .0001$), mPFC (EB: $\chi^2(13) = 130.58$, $p < .0001$; KM: $\chi^2(13) = 62.97$, $p < .0001$), and hippocampus (EB: $\chi^2(13) = 228.68$, $p < .0001$; KM: $\chi^2(13) = 67.00$, $p < .0001$) (Figure S 3). EB-related activity consistently peaked around lag 0, while KM-related peaks varied across regions (0–6s).

These results replicate prior reports of boundary-evoked BOLD activity increases[4,5] and show that KMs also elicit region-specific univariate responses.

.

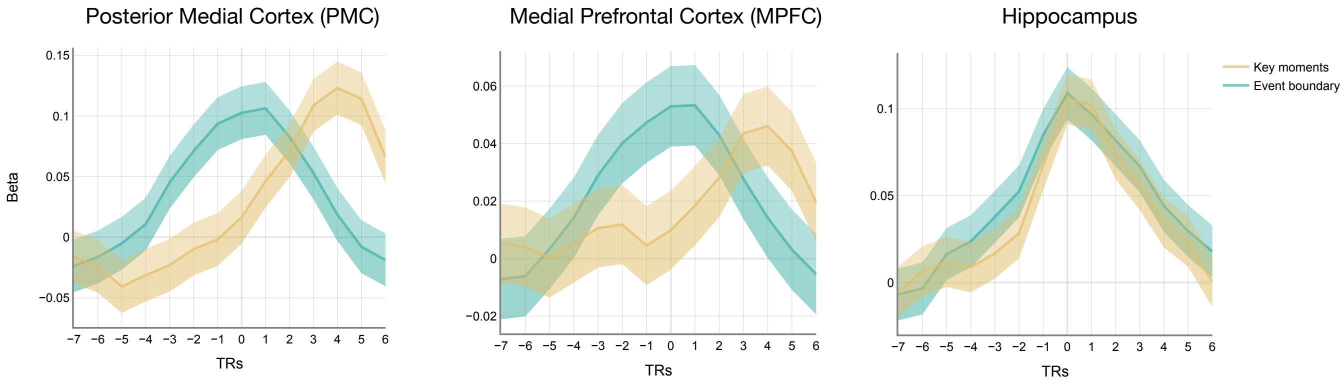


*Figure S 3: Univariate analysis comparing the BOLD activity around event boundaries and key moments in the Posterior Medial Cortex (PMC), Medial Prefrontal Cortex (MPFC), and the Hippocampus (Hipp). Beta values represent the degree of activation across the voxel relative to the baseline and are obtained by the Finite Impulse Response (FIR) model. The solid line represents the linear mixed effects model estimates of beta values. The shaded lines represent the 95% CI.*

### Supplementary analysis S4: The effect of unique key moment and event boundary instants in predicting the ISPS:

We checked whether the unique probability of being a KM or EB predicted the ISPS. The analysis for this followed the same procedures as the ISPS analysis (Figure 3). Except, the predictor variables were the unique KM and unique EB probabilities. The unique probabilities of both the metrics were obtained by regressing out the common variance between the two distributions. A schematic of the analysis is displayed below (Figure S 4). The results replicated the findings of the ISPS analysis (Figure S 5, Table S 1).

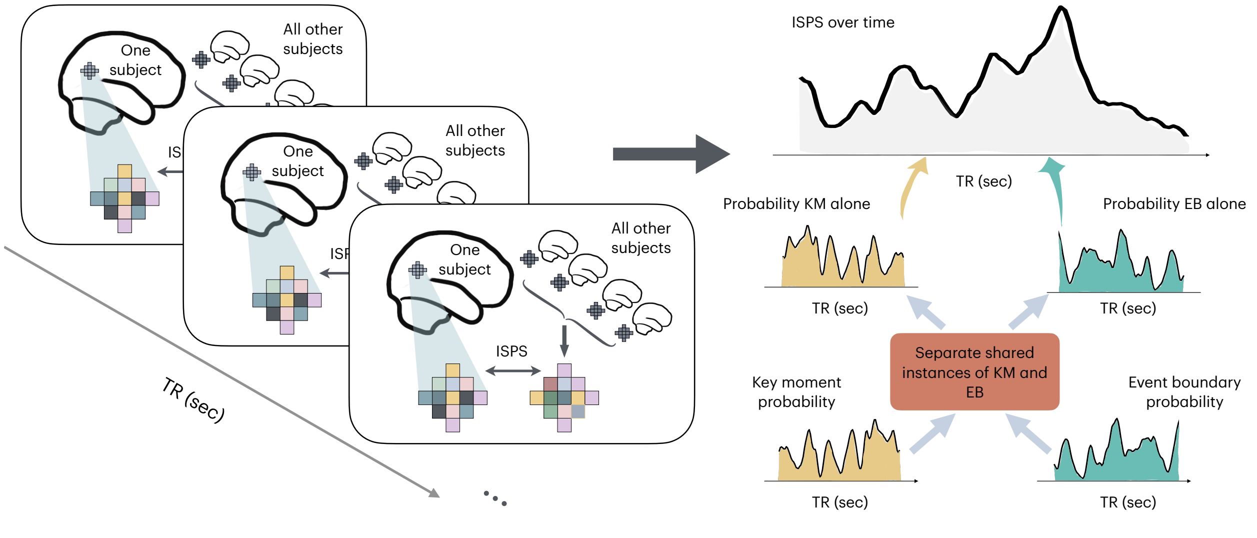


*Figure S 4: Intersubject Neural Pattern Similarity schematic as a function of unique KM and unique EB probability distributions. The unique KM and EB probability distributions were obtained by regressing out the shared components between the two, and the left-over residual from the linear model is used to obtain the unique components of the KM and EB distributions, respectively.*

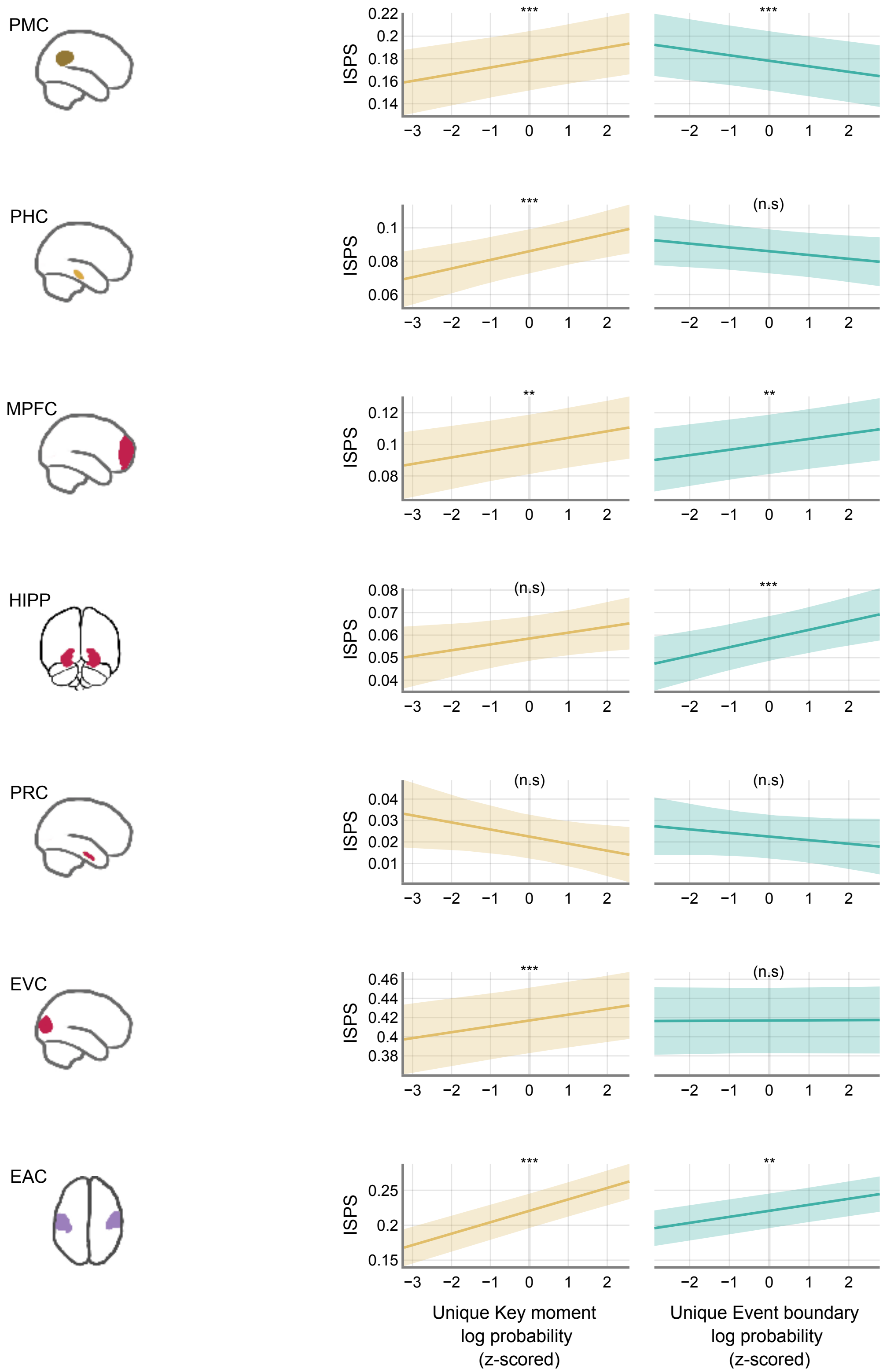


*Figure S 5: Comparing Inter-Subject neural Pattern Similarity (ISPS) against the unique key moment and event boundary probabilities. The first column indicates the ROI that was examined. The second column plots the*

*relationship between the ISPS and the unique key moment probability distribution after log-transformed and z-scored. The third column shows the relationship between ISPS and the unique event boundary probability.*

*Table S 1: Linear mixed-effects models comparing the ISPS activity over time with unique key moment and event boundary probability distributions*

| ROI | Fixed effect | Beta | Std. Error | df | t value | Pr (>\|t\|) | Standard deviation of random effects | |
|---|---|---|---|---|---|---|---|---|
| | | | | | | | clip | sbj |
| PMC | Intercept | 0.18 | 0.013 | 25.83 | 13.298 | < 0.0001 *** | 0.04 | 0.04 |
| | Key Moment (unique) | 0.005 | 0.002 | 2.107e4 | 3.381 | 0.0007 *** | -- | -- |
| | Event boundary (unique) | -0.004 | 0.001 | 3.104e4 | -3.350 | 0.0008 *** | -- | -- |
| | Interaction | -0.003 | 0.001 | 3.130e4 | -2.269 | 0.02 * | -- | -- |
| PHC | Intercept | 0.08 | 0.006 | 35.69 | 12.886 | < 0.0001 *** | 0.02 | 0.02 |
| | Key Moment | 0.005 | 0.001 | 1.296e4 | 3.487 | 0.0004 *** | -- | -- |
| | Event boundary | -0.002 | 0.001 | 3.018e4 | -1.825 | 0.06 . | -- | -- |
| | Interaction | 0.003 | 0.001 | 3.102e4 | 3.429 | 0.0006 *** | -- | -- |
| mPFC | Intercept | 0.1 | 0.009 | 53.88 | 10.428 | < 0.0001 *** | 0.04 | 0.02 |
| | Key Moment | 0.004 | 0.001 | 2.865e4 | 3.012 | 0.002 * | -- | -- |
| | Event boundary | 0.003 | 0.001 | 3.134e4 | 3.024 | 0.002 * | -- | -- |
| | Interaction | -0.0015 | 0.001 | 3.136e4 | -1.529 | 0.126 | -- | -- |
| Hipp | Intercept | 0.05 | 0.005 | 47.91 | 11.605 | 1.6e-15 *** | 0.02 | 0.01 |
| | Key Moment | 0.002 | 0.001 | 1.216e4 | 1.897 | 0.05. | -- | -- |
| | Event boundary | 0.003 | 0.001 | 3.007e4 | 3.353 | 0.0008 ** | -- | -- |
| | Interaction | -0.001 | 0.001 | 3.096e4 | -1.081 | 0.27 | | |
| PRC | Intercept | 0.02 | 0.005 | 29.61 | 4.358 | 0.0001 *** | 0.01 | 0.01 |
| | Key Moment | -0.003 | 0.0017 | 2.996e3 | -1.858 | 0.063 | -- | -- |
| | Event boundary | -0.001 | 0.001 | 2.430e4 | -1.093 | 0.274 | -- | -- |
| | Interaction | 0.0002 | 0.001 | 2.87e4 | 0.144 | 0.88 | -- | -- |
| EVC | Intercept | 0.41 | 0.01 | 32.33 | 24.010 | < 2e-16 *** | 0.06 | 0.05 |
| | Key Moment | 0.006 | 0.002 | 2.848e4 | 3.317 | 0.0009 *** | -- | -- |
| | Event boundary | 0.001 | 0.001 | 3.114e4 | 0.121 | 0.903 | -- | -- |
| | Interaction | 0.005 | 0.001 | 3.117e4 | 3.627 | 0.0002 * | -- | -- |
| EAC | Intercept | 0.20 | 0.01 | 44.01 | 17.651 | < 2e-16 *** | 0.05 | 0.03 |
| | Key Moment | 0.016 | 0.001 | 2.968e4 | 11.479 | < 2e-16 *** | -- | -- |
| | Event boundary | 0.008 | 0.001 | 3.137e4 | 7.388 | 2.22e-13 *** | -- | -- |
| | Interaction | 0.001 | 0.001 | 3.138e4 | 1.038 | 0.299 | -- | -- |
